# Steady state ratios in bipartite population models emerge through obligate cross-feeding

**DOI:** 10.64898/2026.07.29.741403

**Authors:** J. Fabian Pohlkotte, Oliver Ebenhöh, St. Elmo Wilken

## Abstract

Cross-feeding, the phenomenon where one organism consumes metabolites secreted by another, is a ubiquitious phenomenon in microbial communities. Obligate, mutually cross-feeding organisms are a special case of mutualism, where each organism produces a resource that the other organism requires for growth. Many obligate, mutually cross-feeding microbial systems have been studied, including naturally occurring as well as synthetically designed communities. Interestingly, in all these experimental systems it has been observed that the communities approach a fixed biomass ratio, independent of the initial biomass inoculation composition. The repeated reports of stable biomass ratios suggest that such ratios are actually a generic property of mutually obligate cross-feeding systems and that, if the system is stable, such a ratio *must* be approached. As a consequence, it should also be possible to explain and predict this ratio from measurable parameters. Here, we explore mutually obligate cross-feeding systems with mathematical models and show that a stable biomass ratio is a general feature of such systems. Moreover, we show how this ratio can be predicted from key parameters that can be interpreted as the *value* of the resource to the consuming organism and the *cost* of producing a resource to the producing organism.

## 1 Introduction

In nature, microbes live in complex communities. The stability of these communities depends on the metabolic interactions between community members, such as cross-feeding of essential small metabolites that are produced by only a small number of species [1]. It was shown in lab evolution experiments that mutualistic cross-feeding can evolve from competition, which indicates that specialisation, division of labour, and the concomitant mutual dependence presents, at least under some conditions, a selective advantage for the individual species [2]. Obligatory cross-feeding presents the special case of this mutualistic interaction, in which interacting organisms mutually depend on complementary metabolites produced by other organisms, and no organism can grow or survive without the others. Interestingly, when studying obligatory pairwise cross-feeding cocultures in isolation, it has been regularly observed that the organism ratio approaches a fixed value, independent of the initial community composition [3, 4, 5, 6, 7, 8]. Remarkably, all of these systems are quite different from each other in terms of interacting species and exchanged metabolites. These repeated reports of stable organism ratios in obligatory cross-feeding co-cultures raise the question of whether a stable ratio is a phenomenon generally found for these types of interactions, and, if so, which external factors and parameters determine the ratio.

Previous work on this subject suggested that cellular economics, i.e. the *value* of an exchanged metabolite for the receiver and the *cost* for the producer, determine the organism ratio in a predictable fashion [8]. Specifically, motivated by general economic theories [9] applied to biological systems [10], it was hypothesised that an equilibrium ratio is attained when the metabolic burden inflicted by the production of the exchanged metabolites is shared equally among the partners. The metabolic burden was postulated to reflect the amount of ATP that could be produced additionally if that resource was not secreted, but instead metabolised internally. By manipulating the metabolic burden, the authors found that experimentally observed organism ratios largely confirm the theoretical predictions.

In this paper, we go a step further and show that, under some conditions, obligatory cross-feeding co-cultures *necessarily* approach a fixed ratio. For this, we develop mathematical models on cross-feeding systems and show that the fundamental structure of the mathematical equations representing obligatory cross-feeding interactions lead to a fixed organism ratio that is an attractive equilibrium state. As a consequence, under the conditions investigated here, obligatory cross-feeding co-cultures grow towards a ratio that is independent of the initial inoculation ratio. Our approach allows calculating the ratio from the model parameters, and thus provides a conceptual framework to predict the ratios from organism-specific parameters. Comparing highly simplified dynamic models with detailed dynamic flux balance models, in which two genome-scale networks interact, illustrates how these parameters can be determined from experiments, or estimated from the stoichiometries of the metabolic networks.

## 2 Results

We present a series of models of increasing complexity to simulate cross-feeding in a bipartite microbial community. The purpose of these models is to (i) illustrate that pairwise cross-feeding leads to stable organism ratios and (ii) understand which parameters affect this ratio. We present five models of increasing complexity.

1. The simplest model (implicit resource exchange) contains only two dynamic variables, and resource flow is simply assumed to be proportional to the abundance of the cross-feeding partner.
2. The next level of complexity (explicit resource exchange) has four dynamic variables, where both the species abundances and resource concentrations are explicitly described.
3. The cost-benefit model extends the explicit resource exchange model by introducing costs for each unit of a resource that is produced and exported.
4. The chemostat model includes an external resource supply as an additional source of growth limitation, as well as a dilution rate.
5. Finally, we study the behaviour of a dynamic community model, in which the metabolism of each partner is described by a detailed genome-scale metabolic model.

Figure 1 shows an overview of the most relevant processes considered for the five cross-feeding models presented here. The partners *N*_0_ and *N*_1_ exchange two resources (*R*_0_ and *R*_1_) with each other in order to grow. The dynamics of this bipartite system are determined by resource production, secretion, uptake and conversion into biomass.

**Figure 1.**
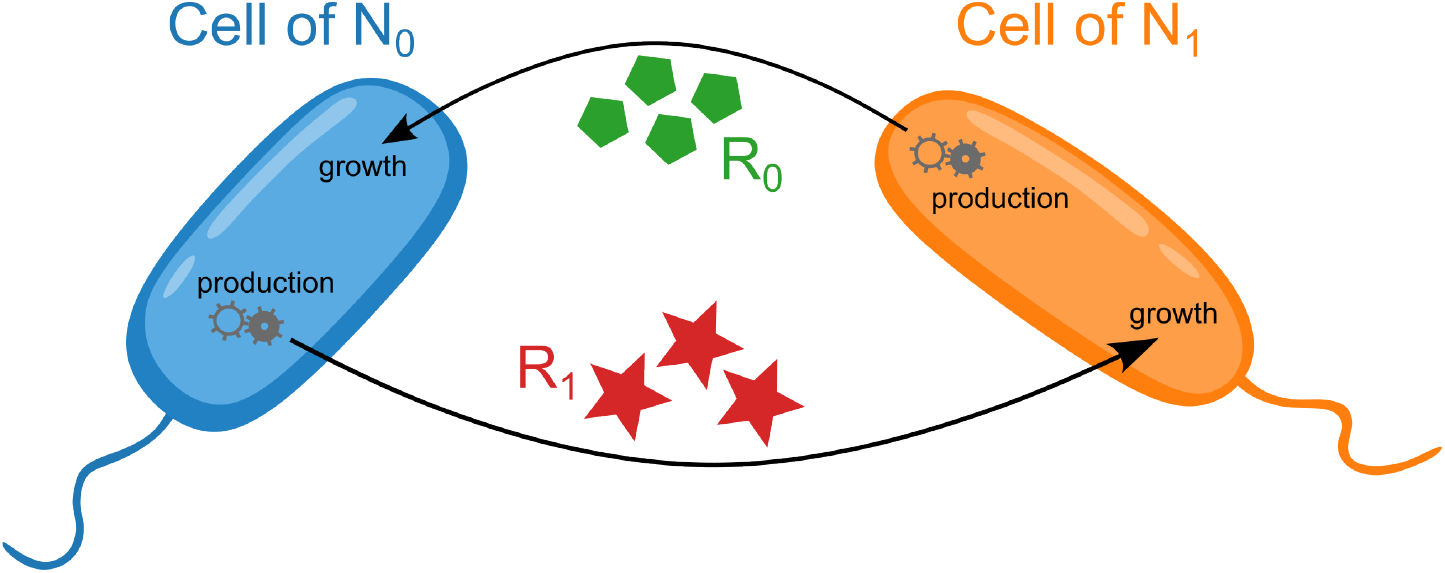
Obligate cross-feeding between two populations. The partners *N*_0_ and *N*_1_ exchange two resources (*R*_0_, produced by *N*_1_ and required by *N*_0_, and *R*_1_, produced by *N*_0_ and required by *N*_1_) with each other in order to grow. The dynamics of this bipartite system are determined by resource production, secretion, uptake and conversion into biomass.

### 2.1 Implicit resource exchange model

Any model describing the growth dynamics of two obligatory cross-feeding microorganisms must reflect this mutual dependency in the equations describing their growth rates. This mutual dependence implies that the growth rate of one partner depends on the presence of the other partner, and *vice versa*.

To construct simplified models of obligatory cross-feeding, we make a number of assumptions. First, we assume that the exchanged resource is the only growth-limiting nutrient, i.e. that all other nutrients, including the carbon and nitrogen sources, as well as macro- and micro-nutrients are present in high abundances. This is a central assumption, which holds for most models presented here. Second, we assume that biomass-specific resource production rates are constant, meaning that the total production rate of the resources is proportional to the biomass of the producing microorganism, i.e. we assume balanced growth.

#### Box 1: Implicit resource exchange model

Growth of the cross-feeding partners *N*_0_ *>* 0 and *N*_1_ *>* 0 is assumed to only depend on the presence of the other partner through the exchanged resource, resulting in the equations

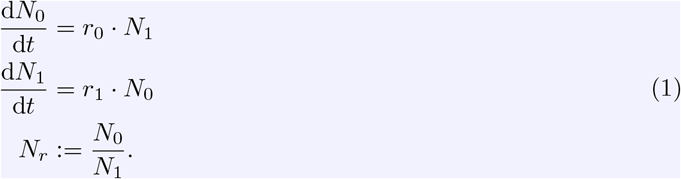

The resources are not modelled explicitly. Instead, the rate parameters *r*_0_ *>* 0 and *r*_1_ *>* 0 represent an assortment of processes: production and secretion of an exchanged resource, its consumption by the partner and the resulting growth. These parameters therefore reflect the costs for the producing partner and the benefits for the consumer of the nutrients. The population ratio *N*_*r*_ of this system converges to

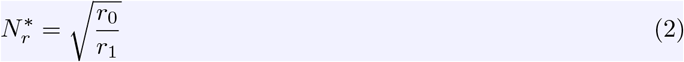

over time for any initial condition (see SI 1). At this equilibrium ratio, both populations grow exponentially with rate

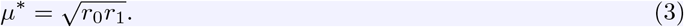

For the simplest possible model, we assume the growth rate to be proportional to the availability of the exchanged resource, and thus proportional to the biomass of the partner organism, leading to Equation (1) in Box 1. The only two variables of the model are the abundances of the cross-feeding partners, *N*_0_ and *N*_1_. The two (positive) specific growth-rate parameters, *r*_0_ and *r*_1_, describe how growth rate of one partner depends on the presence of the other. Thus, these parameters reflect a series of processes, including production and secretion of the exchanged nutrient, its consumption by the partner and the resulting growth. Therefore, these parameters reflect a combination of the *costs* incurred by resource production and the *benefits* experienced by the consumer. It can be shown that the population ratio *N*_*r*_ converges to the equilibrium ratio 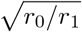 (Equation 2) over time for any initial condition (see SI 1). At this ratio, both populations grow exponentially with the fixed rate 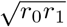 (Equation 3). For this simplest model it can be concluded that a stable abundance ratio must be reached, irrespective of the initial inoculation ratio. The cross-feeding community as a whole exhibits prototophic growth with a specific growth rate that is identical to the geometric mean of the individual specific auxotrophic growth rates.

### 2.2 Explicit resource exchange model

To understand and quantify how different sub-processes affect the auxotrophic growth rate parameters, *r*_*i*_, and thus the stable ratio, we refine our model and explicitly include the exchanged resources as variables. We consider the following processes: production and secretion of resource *R*_0_ (*R*_1_) by organism *N*_1_ (*N*_0_), consumption of *R*_0_ (*R*_1_) by *N*_0_ (*N*_1_), and conversion of *R*_0_ (*R*_1_) into biomass of *N*_0_ (*N*_1_). As before, production and secretion of each resource is assumed to be proportional to the producer’s abundance (new parameters *p*_0_ and *p*_1_). Moreover, the resources are the only growth limiting substrates. We assume that the uptake rate of a resource follows some monotonously increasing function (*c*_*i*_(*R*_*i*_) with *c*_*i*_(0) = 0) of the resource concentration, such as a simple linear relationship or a Monod-like rate law. Finally, we assume the growth yields on the consumed resources to be constant (new parameters *b*_0_ and *b*_1_). This leads to Equation (4) in Box 2.

Like for the simpler implicit resource exchange model (Equation 1), it can be shown that over time, the population ratio converges to an equilibrium value, now given by 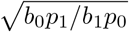 (see Equation 5, SI 2). At this equilibrium ratio, both populations grow exponentially with rate 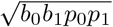 (see Equation 6) and the resource amounts assume a constant non-negative value. Remarkably, these values do not depend on the specific form of the uptake rate functions *c*_*i*_(*R*_*i*_). The only property required for the proof that the equilibrium ratio is unique and stable is that they are monotonously increasing with resource concentration (see SI 2). Once a stable equilibrium ratio is reached and both populations grow with identical rate, the model equations reduce to those of the model with implicit resources (Equation 1), demonstrating that the explicit resources model is a generalization of the implicit resources model.

In Figure 2 we present exemplary simulation results. Panel A illustrates representative time courses of two cross-feeding populations. After an initial phase, both populations grow exponentially with the same growth rate. Panel B demonstrates that a stable biomass ratio is achieved regardless of the initial conditions, approaching the predicted ratio given by Equation (5). In contrast to the populations, the resources converge to a finite value (panel C). The sensitivity analysis presented in panel D shows the dependence of the stable ratio

**Figure 2.**
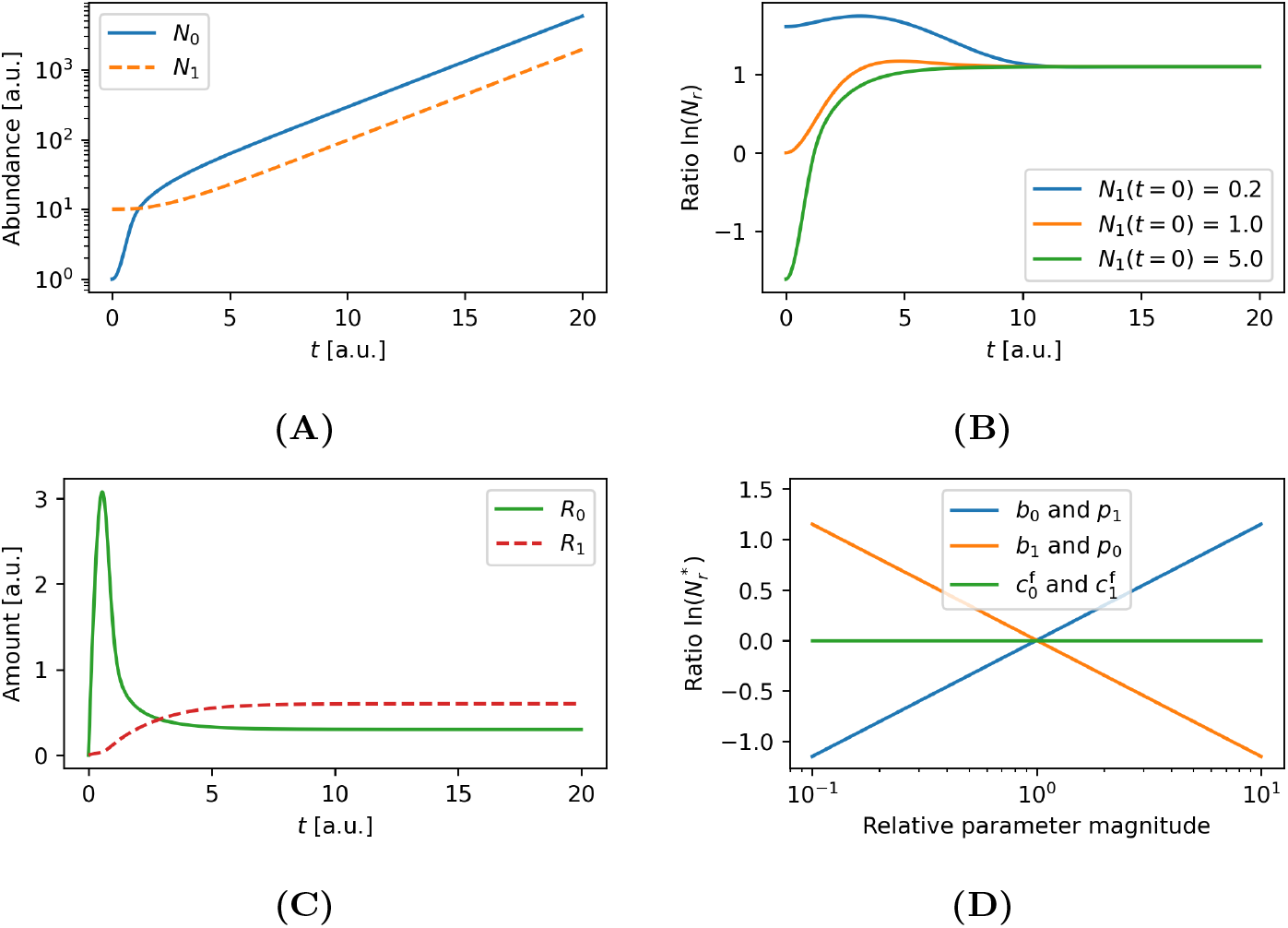
Numeric simulations of the explicit resource exchange model. Shown are simulation results for the explicit resource exchange model described by Equation (4), with resource production rate (*p*_*i*_) and biomass yield (*b*_*i*_) for partner *i*. Here, the linear (unbounded) consumption rate functions 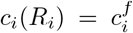.*R*_*i*_ were employed. **(A)** Population abundances over time for 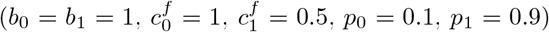After a transitional phase, both populations converge towards a stable ratio and grow exponentially with the same relative rate. **(B)** Population ratios over time for the same parameters as in (A) for different initial abundance ratios. Regardless of the initial condition, the ratio converges to the value predicted by Equation (5). **(C)** Resource amounts over time from the simulation used for (A). The resources stabilize to finite values after a transitional phase. **(D)** Sensitivity analysis of long-term population ratios for different values of the biomass conversion rates *b*_*i*_, the resource production rates *p*_*i*_ and the linear consumption rates 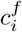.

#### Box 2: Explicit resource exchange model

Two organisms and two resources they exchange between them are considered. *R*_0_ (*R*_1_) is produced by *N*_1_ (*N*_0_), consumed by *N*_0_ (*N*_1_) and converted into biomass of *N*_0_ (*N*_1_), leading to the equations

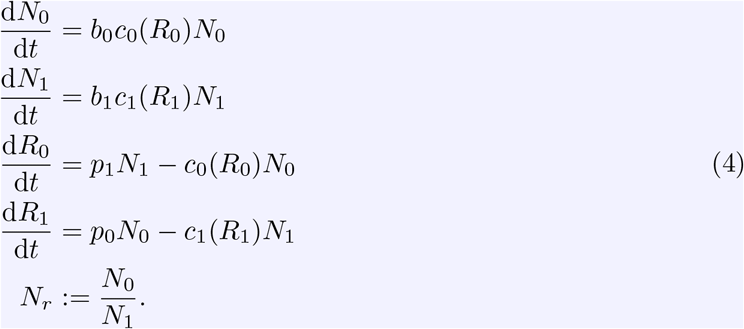

Production and excretion of each resource is assumed to be proportional to the producer’s abundance (parameters *p*_0_ *>* 0 and *p*_1_ *>* 0). The consumption rate functions *c*_0_(*R*_0_) and *c*_1_(*R*_1_) are assumed to be monotonously increasing with the amount of the respective resource and go through the origin. The growth yields on the consumed resources are the constants *b*_0_ *>* 0 and *b*_1_ *>* 0. In the long term, both populations grow exponentially (see SI 2), either one or both resources assume a constant non-zero value and the other resources grows exponentially (if only one resource stabilizes). The population ratio *N*_*r*_ of this system converges to

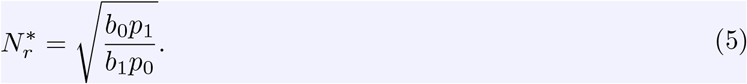

over time for any initial condition if both resources stabilize, which is the most relevant case. At this equilibrium ratio, both populations grow exponentially with rate

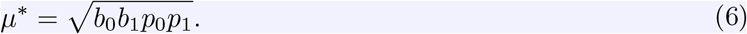

on the model parameters. Interestingly, the stable ratio is not dependent on the functions *c*_*i*_(*R*_*i*_) describing the resource uptake rates (see Equation 5). However, increasing the biomass yield on the exchanged resource (parameters *b*_*i*_) shifts the ratio in favor of that organism. Conversely, increasing the production capacity of the exchanged resource for one organism (parameters *p*_*i*_) will shift the ratio in the opposite direction.

Whereas the population ratio is independent of the specific form of the resource uptake functions, the final resource concentrations are not. For unbounded uptake functions, as chosen for Figure 2, resource concentrations will always converge to a finite value. However, for bounded functions, the situation can occur that resources are produced with a higher rate than the maximal uptake capacity of the cross-feeding partner. This may lead to a situation in which one of the resources accumulates exponentially while the other remains finite (see SI 2.2 for a detailed discussion).

### 2.3 Cost-benefit model

In the explicit resource exchange model (see Equation 4), the parameters *p*_*i*_ denote the biomass-specific nutrient production rates. So far, we assumed that this parameter is a constant. This is realistic if the nutrient is a catabolic by-product, such as ethanol in the case of anaerobic glucose fermentation in yeast [11]. However, in general nutrient production will incur a cost for the producing organism, and resource allocation principles [12, 13] imply that any resource allocated to nutrient formation cannot be used for biomass production and thus growth. As a consequence, increased nutrient production will lead to a reduced growth rate. It therefore appears plausible that this parameter is highly regulated to avoid unnecessary over-production. This trade-off between biomass and nutrient formation has not been explicitly included in the models described in the previous sections.

We extend the explicit resource exchange model to reflect this trade-off by introducing costs for resource production, leading to Equation (7) in Box 3. The new parameters *β*_*i*_ reflect the *costs*, measured by the reduction of growth rate per nutrient produced, incurred by the producing organism. Unlike the explicit resource exchange model, the cost-benefit model is bistable. Depending on the initial conditions, either both populations go extinct, or they grow exponentially at a fixed community growth rate (for details, see SI 3). The dependence of the cost-benefit model on the initial conditions is demonstrated in Figure 3. Here, the linear (unbounded) consumption functions 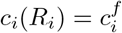.*R*_*i*_ were used. If the resource amounts and population levels are too low initially, both populations will go extinct. At higher initial values, both populations will grow exponentially, reaching the stable equilibrium ratio given by Equation (8).

**Figure 3.**
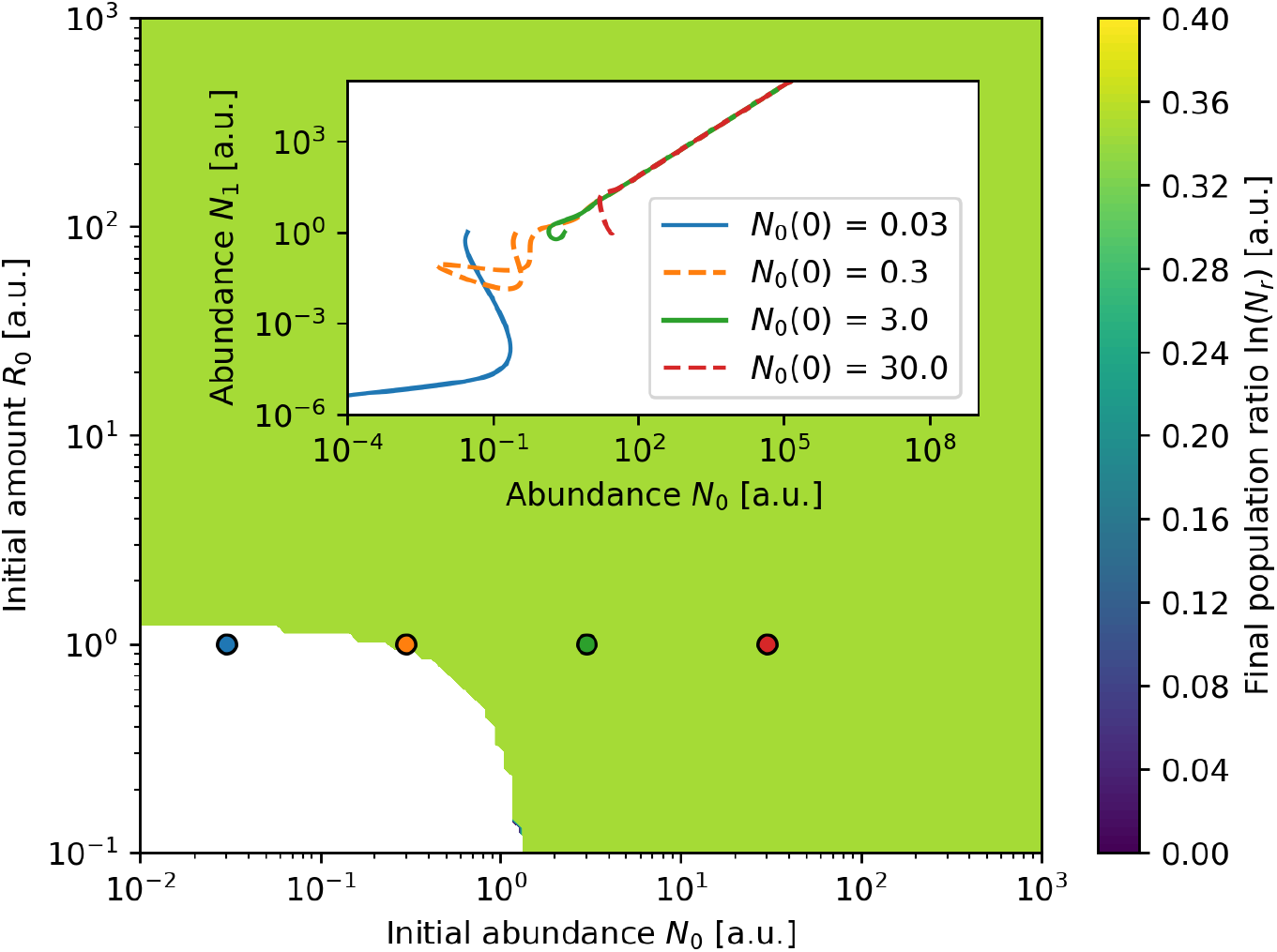
Impact of initial conditions on long-term behaviour. If the initial resource amounts and population levels are too low, both populations will go extinct. When starting at higher initial values, both populations will converge towards the same stable ratio and grow exponentially with the same fixed rate. The ratio and the rate only depend on the parameter values. The linear consumption functions 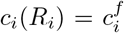.*R*_*i*_ are used. The parameter values are 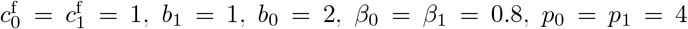 were used. The initial conditions for each simulation were *N*_1_(0) = 1, *R*_1_(0) = 0. **Inset:** Exemplary time series phase diagram illustrating the transitional phase and long-term growth or demise of both populations for the same parameter set and consumption functions. The initial conditions for each simulation were *N*_1_(0) = *R*_0_(0) = 1, *R*_1_(0) = 0 and their starting points are marked on the main figure.

The simulations presented in Figure 3 are exemplary for the case of unbounded resource consumption functions. However, realistically the biomass-specific nutrient uptake rates are always bounded. If, for example, we assume Monod-like uptake rate functions,

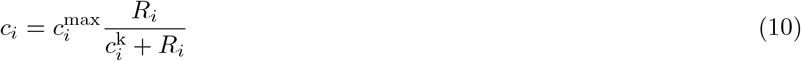

(with half-saturation parameters 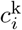 and maximal uptake rates 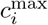), any pa-rameter value 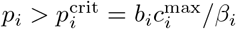 results in d*N*_*i*_*/*d*t <* 0 (see Equation 7) andas a consequence both populations will go extinct. In the case that both parameters *p*_0_ and *p*_1_ are smaller than the critical value, a stable ratio with exponential

#### Box 3: Cost-benefit model

Two organisms and two resources they exchange between them are considered. *R*_0_ (*R*_1_) is produced by *N*_1_ (*N*_0_), consumed by *N*_0_ (*N*_1_) and converted into biomass of *N*_0_ (*N*_1_), leading to the equations

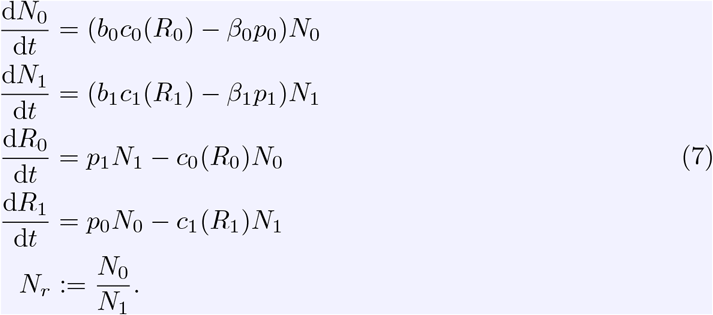

Production and secretion of each resource is assumed to be proportional to the producer’s abundance (parameters *p*_0_ *>* 0 and *p*_1_ *>* 0). Resource production reduces an organism’s growth rate as a cost (parameters *β*_0_ ≥ 0 and *β*_1_ ≥ 0). The consumption rate functions *c*_0_(*R*_0_) and *c*_1_(*R*_1_) are assumed to be monotonously increasing with the amount of the respective resource and go through the origin. The growth yields on the consumed resources are the constants *b*_0_ *>* 0 and *b*_1_ *>* 0.

For some initial conditions, both populations go extinct (see SI 2). In all other cases, both populations grow exponentially in the long term, either one or both resources assume a constant non-zero value and the other resources grows exponentially (if only one resource stabilizes). The population ratio *N*_*r*_ of this system converges to

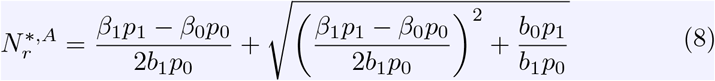

over time if both resources stabilize, which is the most relevant case. At this equilibrium ratio, both populations grow exponentially with rate

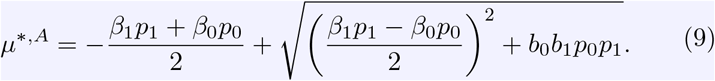

growth is an attracting state (see SI 3 for a mathematical proof). However, the growth rate and the stable abundance ratio now depend on whether both resource concentrations will remain finite or whether one of the resources increases indefinitely. Figure 4 exemplifies this dependence of the stable ratio (panel A) and the growth rate (panel B) on the parameters *p*_*i*_. The analytic formulas for stable abundance ratios and corresponding growth rates in each case are derived in SI 3. Interestingly, maximal growth rate is achieved exactly at the bifurcation point between the three possible modes of system behaviour. At this point, increasing the production rate of one resource will result in an over-production and thus to an accumulation of that resource. Thus, the optimal population growth rate is found at the parameter combination at which the consumption capabilities of both populations are fully exploited without overproducing either resource.

**Figure 4.**
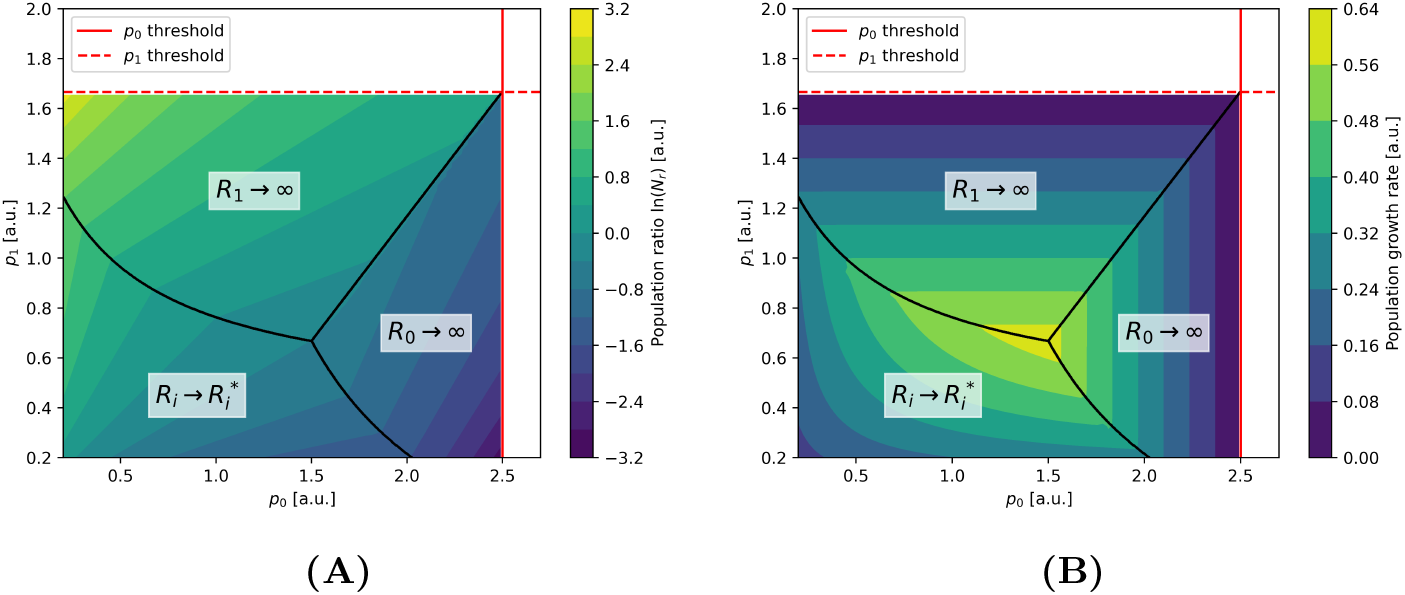
Steady-state abundance ratios and growth rates for different production rates. Increasing either *p*_*i*_ beyond their respective thresholds 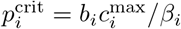 causes both populations to go extinct for all initial conditions (blank space). Below that threshold, high enough starting populations and resource levels lead to both populations growing exponentially at a fixed ratio. Stable ratio **(A)** and community growth rate **(B)** depend on the production rates *p*_0_ and *p*_1_. For lower production rates (left lower section in the graphs), the resource amounts stabilize at a finite value and the population growth rate is given by Equation (9). For higher production rates, only one resource stabilizes, while the other grows exponentially. Here, the growth rate linearly decreases with *p*_0_ for *R*_0_ → ∞ and with *p*_1_ for *R*_1_ → ∞ . The consumption functions, Equation (10), and parameter values 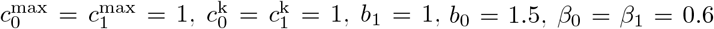 were used. The initial conditions for each simulation were *N*_*i*_(0) = 1, *R*_*j*_(0) = 200. For the chosen parameter values, this ensures that the stable equilibrium ratio will be reached if it exists.

**Figure S1.**
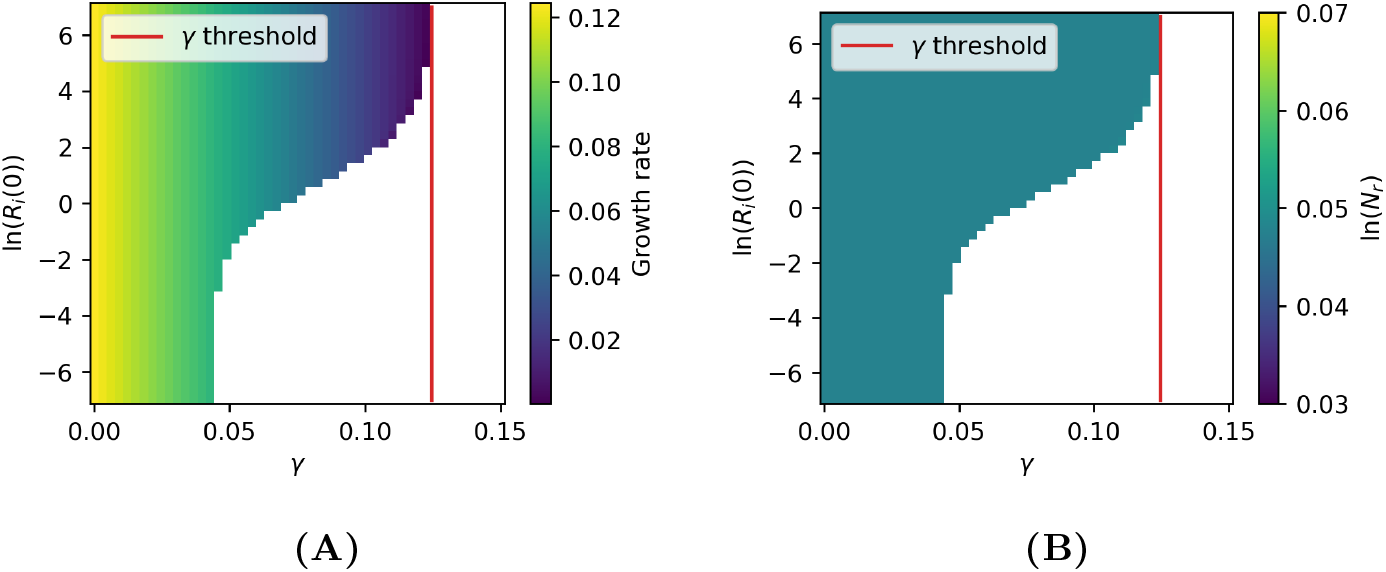
Steady-state growth rates and population ratios for different dilution rates and initial conditions. The populations go extinct if the initial resource levels are too low (blank space). This minimum requirement rises with increasing dilution rate *γ*. There exists a threshold for *γ* beyond which no initial conditions can lead to growth. **(A)** Long-term population growth rates are exactly the intrinsic growth rates (with no dilution, *γ* = 0) minus the dilution rate. **(B)** Long-term population ratios – the same value is reached regardless of *γ* and the choice of initial conditions, as long as the populations do not go extinct. The consumption functions 10 and parameter values 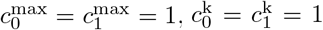, *b*_1_ = 1, *b*_0_ = 1.1, *β*_0_ = *β*_1_ = 0.8, *p*_0_ = *p*_1_ = 0.5 were used. The initial conditions for each simulation were 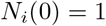.

### 2.4 Chemostat conditions

In realistic conditions, dilution, spatial displacement, predation by other populations and metabolite degradation can constantly reduce nutrient concentrations and organism abundances. We therefore investigate how the system behaviour is affected if we assume a constant dilution (rate *γ*), resulting in Equation (S33). Remarkably, introducing a constant dilution does not alter the fundamental system behaviour. It can be shown (see SI 4.1) that also with a constant dilution rate the system is bistable with either extinction (*N*_*i*_ = 0), or exponential growth with a fixed organism ratio as the two attracting states. In fact, for the

#### Box 4: Other growth-limiting nutrients

This model in an extension of the cost-benefit model (Equation 7). Adding a dilution rate *γ* and an externally provided resource which may be growth limiting leads to the equations

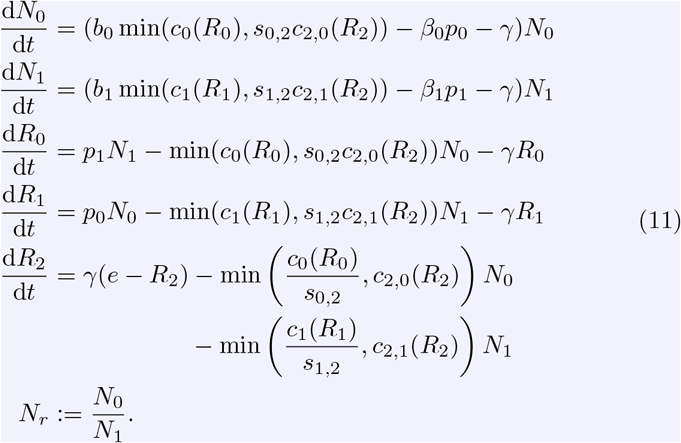

Both populations and all three resources are diluted with rate *γ. R*_2_ increases with the absolute rate *γ* · *e*, where *e* denotes the concentration of the external resource in the feed. Both partners require this resource for biomass production which can now be limited either by the externally supplied resource or one of the exchanged resources. *s*_0,2_ and *s*_1,2_ describe the amount of *R*_0_ and *R*_1_ required to convert one unit of *R*_2_ into biomass of *N*_0_ or *N*_1_.

stable exponential growth state, the overall growth rate is reduced exactly by the dilution rate *γ* when compared to the model without dilution, while the organism ratio is unaffected. Moreover, the basin of attraction for the extinction state is enlarged with increasing dilution rate (see Figure S1).

### 2.5 Other growth-limiting nutrients

Every model discussed so far is based on the assumption that the only growth limiting nutrient is the exchanged resource. This is realistic if all other nutrients are present in high abundance. In a batch culture, this condition can only last temporarily, because eventually one of the other nutrients will be depleted to such an extent that it becomes growth limiting. A typical example is a coculture of two bacterial amino acid auxotrophs cross-feeding complementary amino acids, while both species consume glucose as a carbon source.

Because stable organism ratios are also observed in batch cultures (or repeated batch growth with transfers to fresh medium) [7], we investigate how well our idealised models can predict the final organism ratio in a co-culture. For this, we include a third dynamic resource, *R*_2_, that both organisms require to grow. Now, the growth of each organism can be limited by one of two resources, and, following the arguments provided in [14], we simply assume that the relative growth rate is determined by the minimum allowed by each resource individually, i.e.

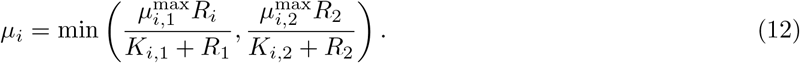

The full formulation of the model is given in Equation (11), Box 4.

Figure S2 illustrates the dynamics of this model. Here, we assumed Monod-type resource uptake functions (see Equation 10) and no dilution (batch conditions, *γ* = 0). Initially, both populations grow approximately exponentially. In this phase, the exchanged nutrients are growth limiting and, as in the models discussed above, the abundances converge towards a stable ratio. However, when the external resource *R*_2_ becomes limiting, growth rate is reduced and approaches zero. Clearly, the longer the exponential phase lasts, the closer the abundances will approach the theoretical value predicted from the simplified models without external resource. This is illustrated in Figure 5, where the final ratios are plotted against the initial external resource concentration. While low concentrations result in a short exponential growth phase and thus in organism ratios rather different from the value expected by the simplified models, for high initial concentrations the achieved ratio converges to that predicted by the simpler models.

**Figure 5.**
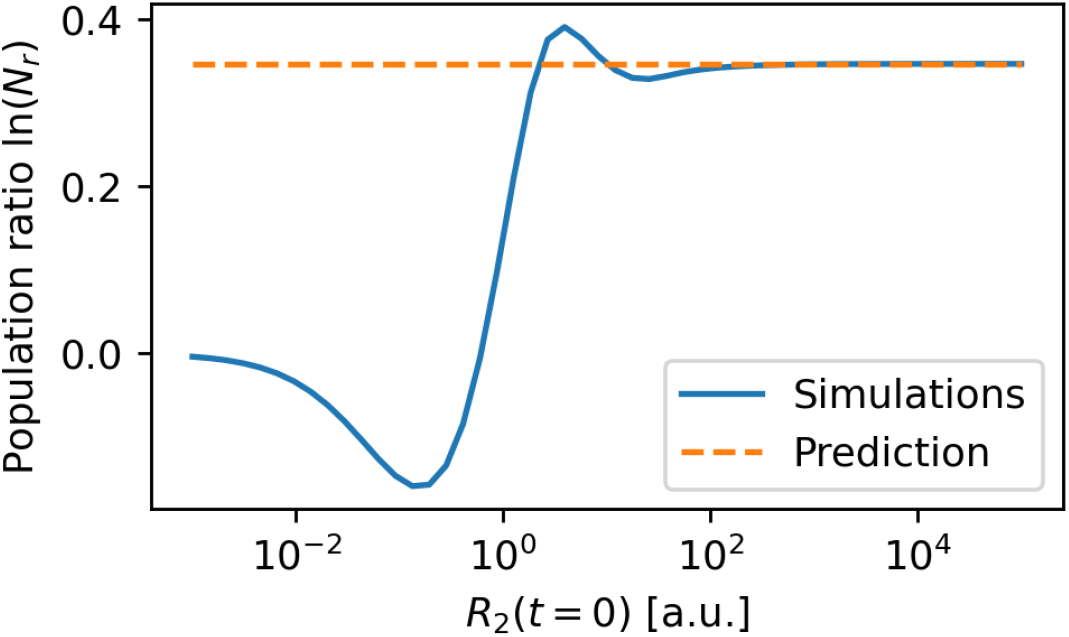
Approximation of exponential growth dynamics in batch cultures. The external resource model (Equation 11) without dilution (*γ* = 0) was used to simulate batch culture conditions. The Monod consumption functions with upper limits were used (Equation 10). The parameters were set to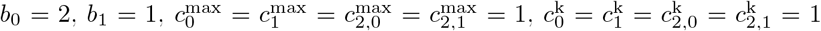 *β*_0_ = *β*_1_ = *γ* = *e* = 0, *p*_0_ = *p*_1_ = 0.5, *s*_0,2_ = 1 and *s*_1,2_ = 10. Shown are the final population ratios in a simulated batch experiment for different initial values of the external resource *R*_2_. For lower values, the equilibrium ratio obtained from the cost-benefit model (Equation 8) differs greatly from the simulation results. However, high enough starting amounts lead to agreement between simulations and the analytical prediction.

### 2.6 Obligatory cross-feeding in coupled genome-scale networks

Lastly, we test whether our analytic results derived for the simplified models also hold if the metabolism of the cross-feeding organisms are described by more complex models. We constructed three variants of a genome-scale metabolic model of *E. coli* (iJO1366 [15]), which have been made auxotrophic for a single amino acid (isoleucine ΔI, lysine ΔK, and threonine ΔT) by removing single reactions in their respective biosynthesis pathways. To simulate the cross-feeding pairs, export reactions of the corresponding amino-acids were added, resulting in pairs of models that can only produce biomass if they cross-feed complementary amino acids with their partner. We studied the dynamics of these obligatory cross-feeding genome-scale networks by simulating the population dynamics using dynamic Flux Balance Analysis (dFBA) [16], implemented in the COBRApy software package [17]. All three possible obligate cross-feeding pairs (ΔI-ΔK, ΔK-ΔT and ΔI-ΔT) investigated here displayed the same qualitative dynamic behaviour.

Figure 6 exemplifies the results for the ΔI-ΔK cross-feeding pair. After an initial transitory phase, both strains approach exponential growth (panel A), regardless of the initial population ratio (panel B). Whereas the populations grow exponentially, the extracellular resource concentrations approach a constant finite value (panel C). In order to test whether the simple explicit resource exchange model is able to quantitatively predict the community growth rate (Equations 5) and stable organism ratio (Equation 6), we need to obtain the corresponding parameters *b*_*i*_ (yield) and *p*_*i*_ (biomass-specific production rate) from the genome-scale network models. The parameters *p*_*i*_ are directly defined in the genome-scale models as flux constraints, fixing the biomass-specific secretion rates, while the parameters *b*_*i*_ were obtained from the definition of the biomass reaction in the iJO1366 model. Explicitly, the molar fraction of the respective amino acid in the biomass reaction is the inverse of the biomass yield on the amino acid, and was used to parameterize *b*_*i*_. These parameters result in a precise prediction of the exponential growth rate and the stable ratio (see red dot in panel C of Figure 6).

**Figure 6.**
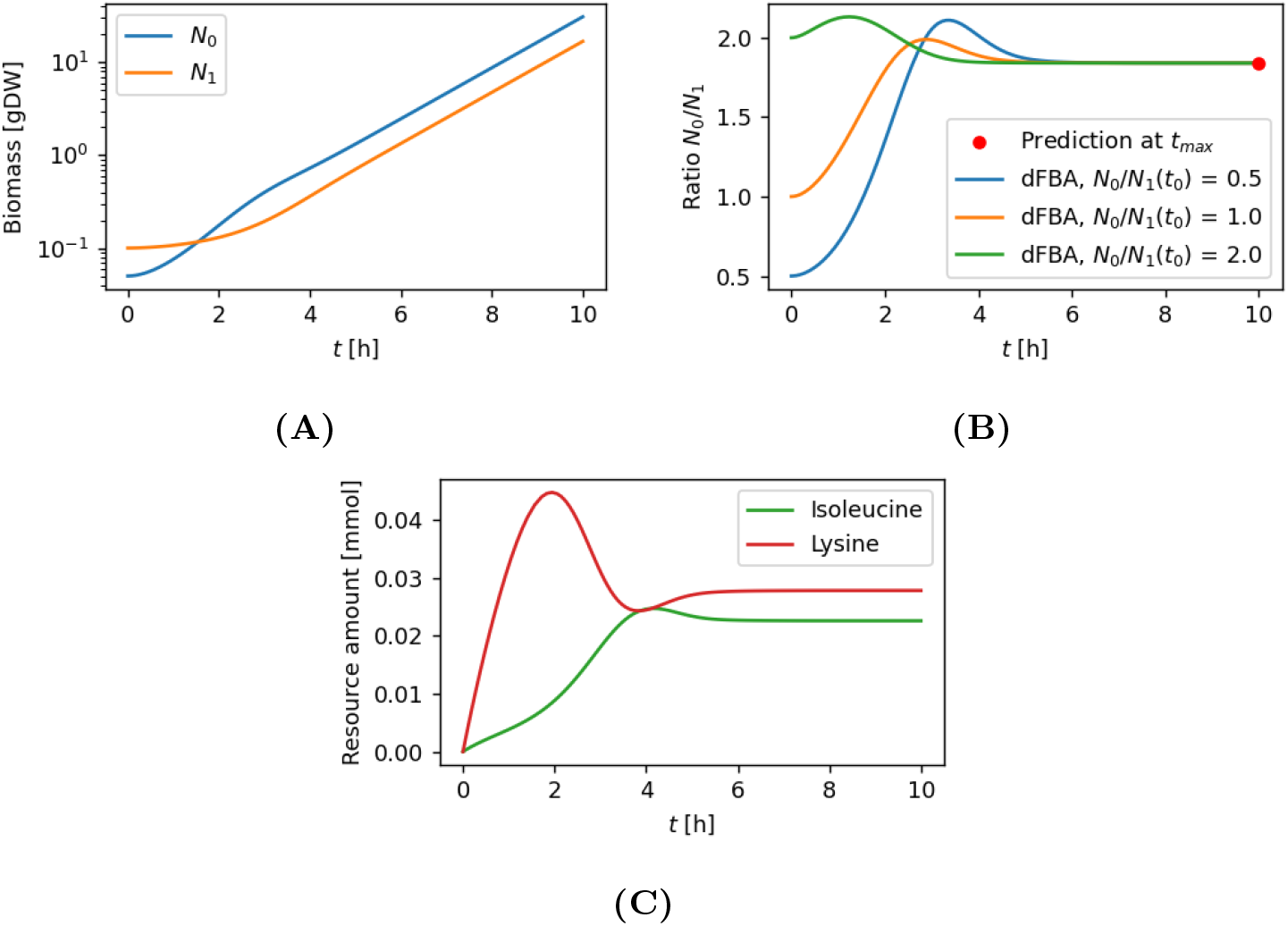
Simulation of two coupled genome-scale networks with obligatory cross-feeding of the amino acids isoleucine and lysine. Dynamic FBA was applied to simulate the dynamics of two genome-scale metabolic networks of *E. coli*, each of which was modified to become auxotroph on one of the animo acids isoleucine (ΔI) or lysine (ΔK). *N*_0_ is the isoleucine producer, *N*_1_ the lysine producer. The amino acid excretion rates were both set to 0.1 mmol/[gDW·h]. The uptake functions are given by Equation (10). The parameters are set to 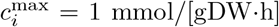 and 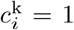. Both populations can grow due to the exchange of the amino acids. **(A)** Population abundances over time. After a transitional phase, both populations grow exponentially with the same rate and a constant ratio *N*_0_*/N*_1_. **(B)** Predicted final population ratio (red dot) and simulated ratios over time for different starting ratios. After a lag phase, the simulated ratios all stabilize at the same level. The composition prediction was calculated using Equation (5) of the explicit resource exchange model (Equation 4). Remarkably, simulation and prediction are in agreement. **(C)** Amino acid amounts over time for the simulation shown in (A). After the transitional phase, both resource levels stabilize.

The predictions also exactly match in the case of the ΔK-ΔT obligate cross-feeding pair (Figure S3). However, applying Equation (5) to predict the stable organism ratio for the ΔI-ΔT cross-feeding pair (Figure S4) results in a quantitatively wrong result. The reason for this discrepancy is that in this scenario a fundamental assumption of the explicit resource model (Equation 4) is violated. The resource production pathways for isoleucine and threonine are not independent. In fact, threonine is a precursor for isoleucine production and therefore, the isoleucine production rate of the ΔT auxotroph is dependent on threonine supply by the ΔI auxotroph. Despite this circular dependency, an analytical formula for the stable organism ratio can also be derived for this scenario (see Equation S35 and SI 5 for the derivation). This further underscores the result that stable ratios are an inherent property of mutually obligate cross-feeding communities.

## 3 Discussion

Both the implicit and explicit resource exchange models offer simple cross-feeding mechanisms by which population abundances can reach stable equilibrium ratios. The explicit resource exchange model predicts the same ratio as metabolic burden theory [8], but through a different approach. This suggests that the overall mechanism that leads to a stable ratio is a fundamental result of this type of resource exchange-based interdependence of two populations regardless of the differences in ATP generation potential, or of resource production capabilities. For the explicit resource exchange model with unbounded resource consumption functions, only the parameters denoting resource-to-biomass conversion and resource production capabilities are relevant for the long-term behaviour of the co-culture.

Interestingly, the equilibrium ratio is not a function of the resource consumption capabilities of the cross feeding community. This suggests that the resource production and biomass production represent the bottleneck for resource exchange processes in microbes, while the resource consumption rate might only play a minor role in determining the stable ratio. However, when one or both of the resource consumption functions have an upper limit, the resulting equilibrium ratio may depend on maximum consumption capabilities. Therefore, the resource consumption processes can be relevant whenever interactions between microbes tend to involve resource concentrations that nearly or completely saturate the resource uptake of those consumption processes. For these cases, the explicit resource exchange model predicts the possibility of one resource to grow towards infinity. However, this is not expected to be reflected in nature because production processes can be down-regulated when a metabolite concentration is very high in order to avoid wasting energy and resources.

One of the core assumptions of all models presented here is that all resources are unlimited except for the ones that are being exchanged between populations. Likewise, we assume that resource production is constant, and not subject to regulation (see [18] for an elaboration) or mutational pressures to escape the community constraints. Additionally, space is also assumed to be unlimited.

Despite this, the results are still relevant for different cases, both in the laboratory as well as in nature. Experiments conducted in a chemostat effectively create these conditions of infinite space and resources through continuous input of fresh medium and dilution. For batch cultures, space and resources are limited only after a significant biomass amount or resource depletion have been reached. Therefore, any setup with a sufficiently high initial resource concentration and a sufficiently low initial biomass will match the model assumptions for an intermediate time span during which a stable population ratio can be reached.

The models we presented here place restrictions on what types of resources each population produces, as well as the quantity and direction of the exchange. These restrictions directly give rise to the characteristic stable ratio reached by the models. Tasoff et al. [10] provide a different approach for resource exchange in microbial co-cultures. Their approach does not have the same constraints for resource production and exchange, but also results in a stable equilibrium ratio by allowing the populations to optimize how they allocate production capabilities. This suggests that stable population ratios in microbial communities that interact through resource exchange can arise via different mechanisms. Beyond auxotrophy and simple carbon cascades, stable community compositions have also been observed between microbes coupled through direct electron transfer mechanisms, demonstrating the wide-spread applicability of this theory [19]. In addition, the dFBA simulations for the isoleucine and threonine knockout mutant combination gave rise to a stable equilibrium ratio as well, but the precise mechanism differs from the mechanism implied by the explicit resources model due to the dependence of isoleucine production on threonine. This highlights the limitations of the simplified model since it assumes no such circular dependencies exist for the resource production pathways. Nevertheless, a stable ratio was still observed, reinforcing the argument presented here: an attractive, unique equilibrium composition is a necessary consequence of a growing, mutually obligate community.

## SI 1 Implicit resource exchange model

This section of the supplementary material supports section 2.1 and relates to model 1.

After calculating the derivative of *N*_*r*_, the equilibrium and its stability can be determined.

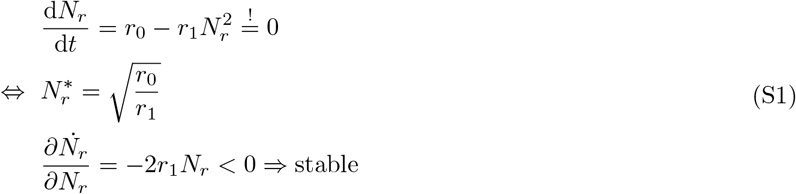

The growth rate at ratio equilibrium *µ*^∗^ is as follows:

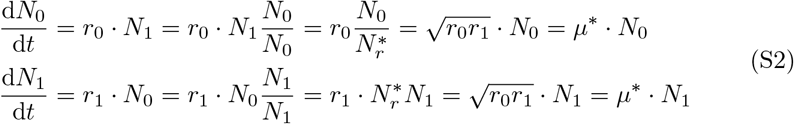

The model can be parameterized easily when both the equilibrium ratio 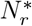 and the equilibrium growth rate *µ*^∗^ are known:

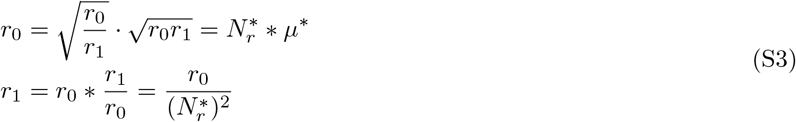

## SI 2 Explicit resource exchange model

This section of the supplementary material supports section 2.2 and relates to model (4).

### SI 2.1 Stability of the equilibrium ratio

Consider the system (*N*_*r*_, *R*_0_, *R*_1_):

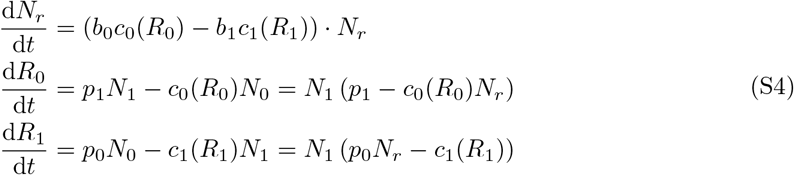

There is only one possible equilibrium

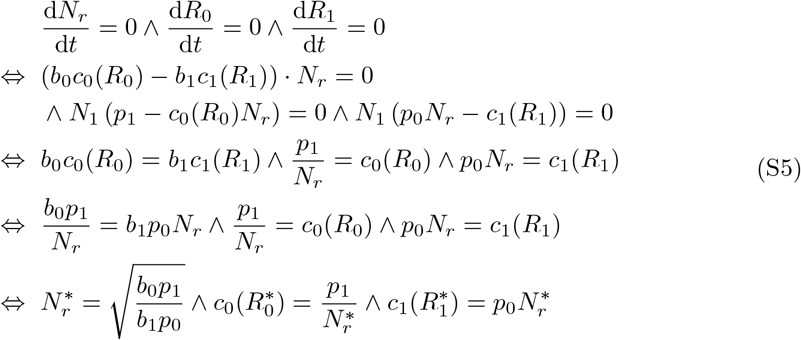

If the resource uptake functions *c*_*i*_(*R*_*i*_) are unbounded, this equilibrium point always exists, independent on the values of the parameters *b*_*i*_ and *p*_*i*_. However, for bounded uptake functions, upper limits for the parameters *p*_*i*_ exist, above which no solution exists. These cases are investigated in detail below in Section SI 2.2.

The Jacobian of the system is:

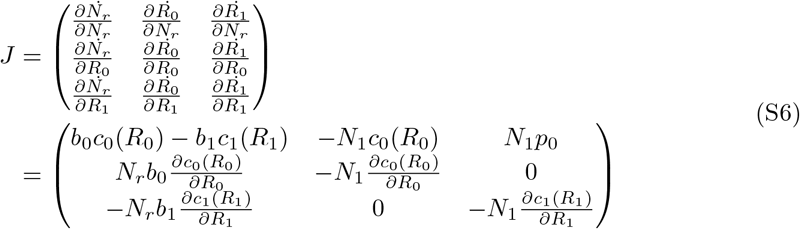

Evaluating *J* at the equilibrium point yields:

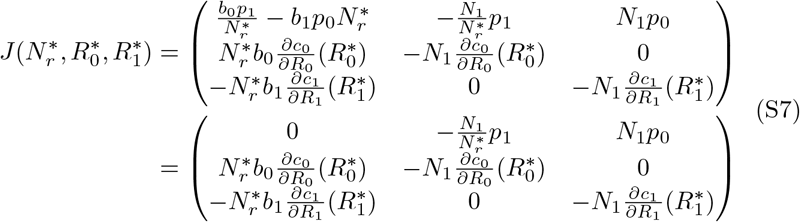

Finding the roots of the characteristic polynomial for *J* ^∗^ yields the eigenvalues:

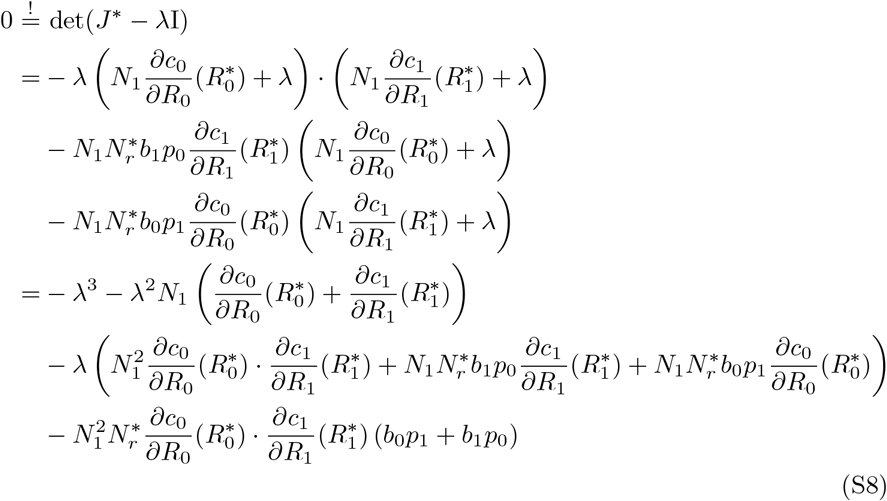

Multiplying the equation above with minus one and defining labels *x*_0_ *>* 0, *x*_1_ *>* 0, *x*_2_ *>* 0 for the coefficients yields a simplified equation. The coefficients are strictly positive because all of their components are as well. Splitting *λ* = *r* +*mi* into its real and imaginary parts results in:

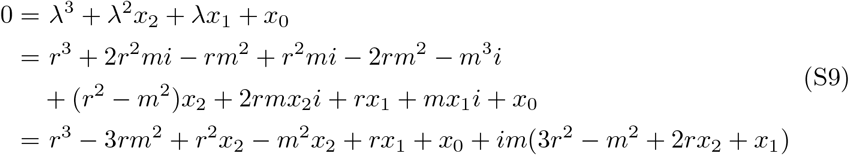

For the polynomial to become zero, both the real and the imaginary part need to be zero:

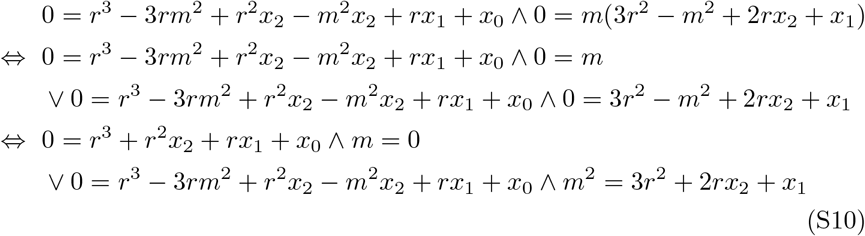

For *m* = 0 it can be seen that the left condition can only be valid for *r <* 0. Inserting *m*^2^ into the left condition for *m*≠ 0 yields:

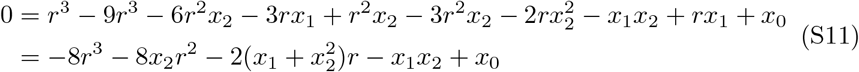

It can be seen that the condition is only valid for *r <* 0, if *x*_1_*x*_2_ *> x*_0_. The latter is proven as follows:

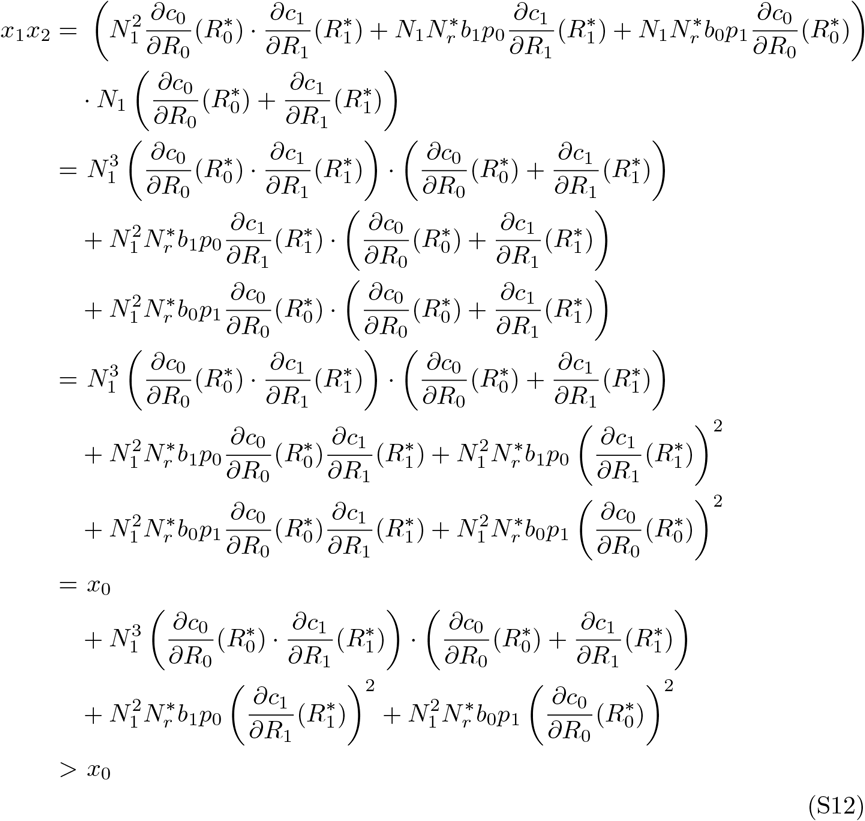

Thus, it is proven that the real part *r* of all eigenvalues has to be negative. This means that the equilibrium is asymptotically stable.

Evaluating the differential equations at ratio equilibrium reveals that they resemble the simple model (1):

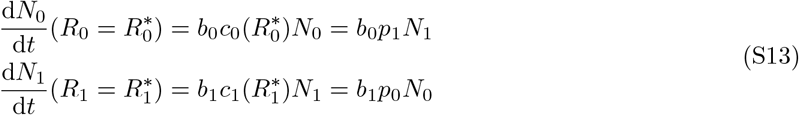

The per-capita growth rate of both populations at ratio equilibrium *µ*^∗^ is the following:

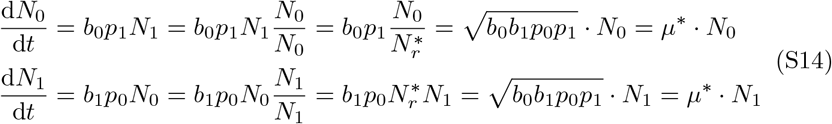

### SI 2.2 Limited uptake kinetics

For cases where one or both resource consumption functions *c*_0_(*R*_0_), *c*_1_(*R*_1_) have an upper limit 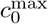 or 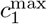 (as is the case for Monod saturation curves, for example), the resource equilibria might not always exist. The existence of them can be determined by evaluating the resource ODEs for a specific population ratio:

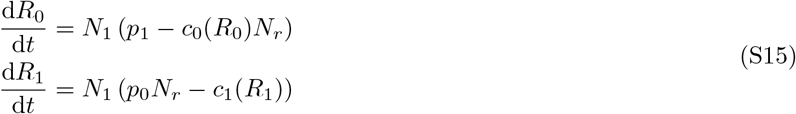

Suppose both consumption functions have an upper limit and the initial population ratio and parameters are such that the following holds:

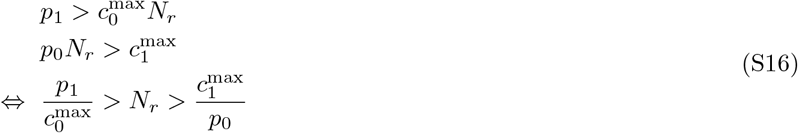

Then, both resource amounts will increase over time, meaning that *c*_0_(*R*_0_) and *c*_1_(*R*_1_) will reach 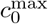 and 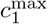 respectively. Applying this to the population ODEs yields:

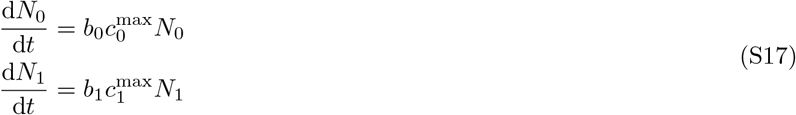

Therefore, both populations will increase exponentially over time with different rates. Suppose 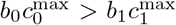 (the other case is analogous). Then, the ratio *N*_*r*_ will increase over time. Considering the resource ODEs again reveals that one of the resource equilibria will stay non-existent (in this case, *R*_1_) whereas the other one will come into existence (here, *R*_0_). Therefore, any scenario starting with both resource equilibria not existing will eventually change towards one equilibrium existing.

Suppose the equilibrium for *R*_0_ exists and the one for *R*_1_ does not exist (the opposite case is analogous). Then, *R*_0_ will eventually reach its equilibrium where *p*_1_*N*_1_ = *c*_0_(*R*_0_)*N*_0_ holds. *R*_1_ will increase, which eventually leads to 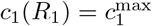. Applying this to the population ODEs yields:

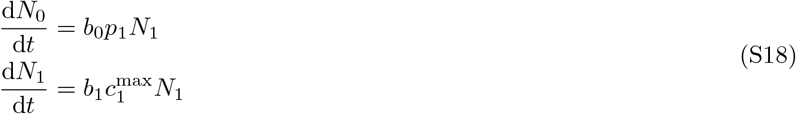

Now, both populations increase with absolute amounts that have a fixed ratio between them. Therefore, both populations will trend over time towards that ratio: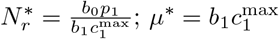. Since any case where both resource equilibria do not exist will revert to a case where one equilibrium exists again, there are only three cases left:

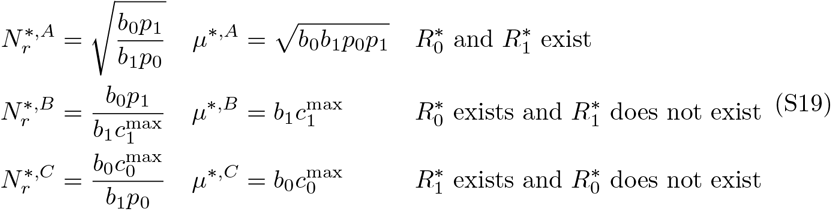

These cases can be differentiated by examining the existence of partial equilibria of both resources. The resource ODE values are evaluated at the upper resource consumption limits 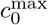 and 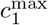 for any one of the three 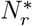 values. Assuming both populations to stay at the same ratio, the partial resource equilibria 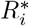 exist if their differential equations at 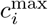 are negative:

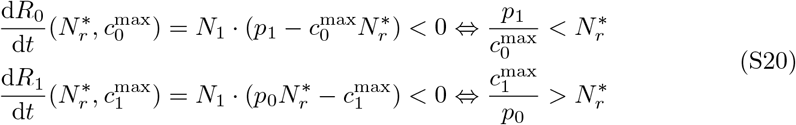

Since each of the possible values for 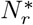 was calculated under the assumption of one of the three scenarios, this calculation can be performed as a test of those assumptions. If there is a contradiction, the scenario can be ruled out. If two tests fail and one succeeds, the 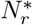 value of the scenario without contradiction will be the steady state ratio of the system.

It can be shown that all transitions between the conditions make one of the three scenarios possible and the other ones impossible, meaning that this proof by contradiction always works: 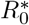 does not exist if 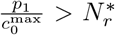. Crossing this threshold through changing a parameter is equivalent to changing between the state 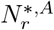 where both resource equilibria exist and the state 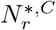 where only 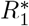 exists:

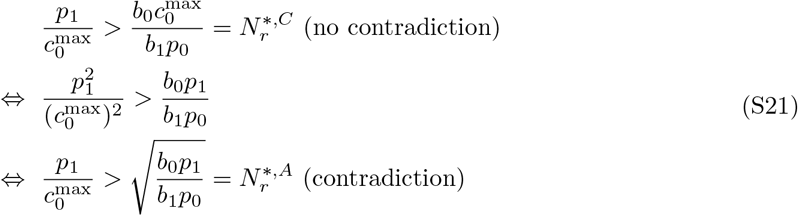

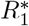 does not exist if 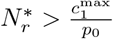. The same kind of state transition can be shown for this condition as well:

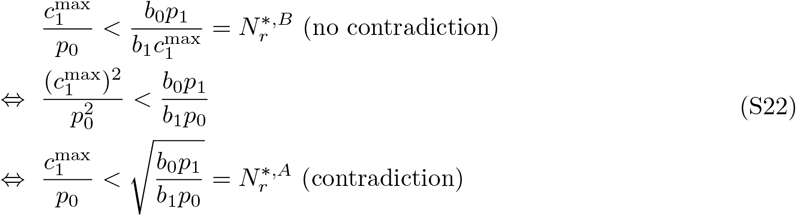

Additionally, a transition exists between 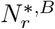 and 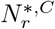:

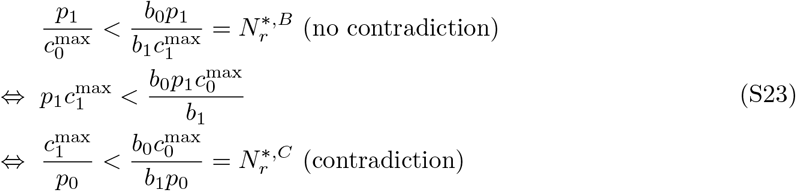

Suppose that 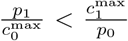. Table S1 shows an overview of the ratio equilibria and their transitions for this case. Now, 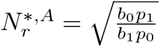 is without contradiction for 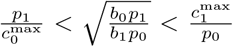 and has a contradiction if it is outside that interval. Violating the lower boundary causes 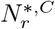 to have no contradiction; violating the upper boundary causes 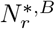 to have no contradiction.

**Table S1.**
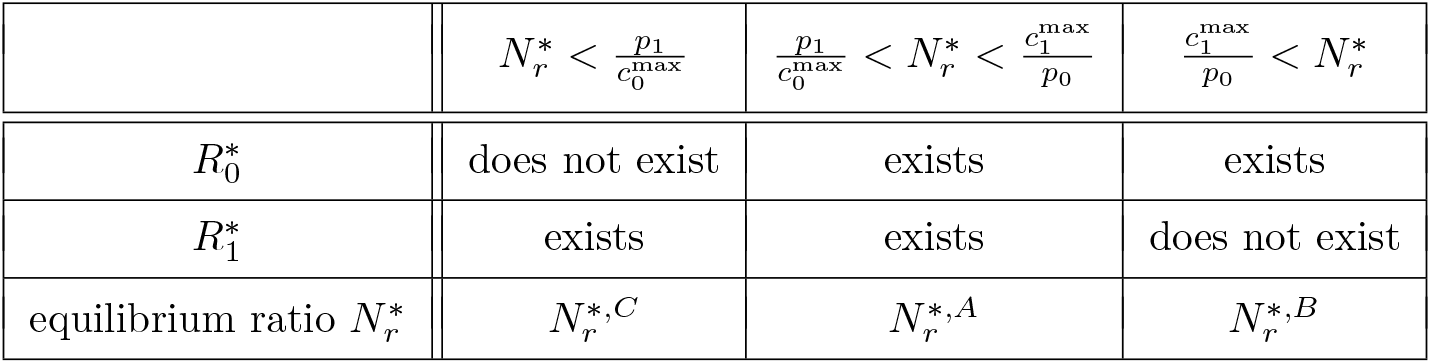
Existence of the resource equilibria and contradictionless equilibrium ratios for 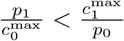.

| | $N_r^* < \frac{p_1}{c_0^{\max}}$ | $\frac{p_1}{c_0^{\max}} < N_r^* < \frac{c_1^{\max}}{p_0}$ | $\frac{c_1^{\max}}{p_0} < N_r^*$ |
| --- | --- | --- | --- |
| $R_0^*$ | does not exist | exists | exists |
| $R_1^*$ | exists | exists | does not exist |
| equilibrium ratio $N_r^*$ | $N_r^{*,C}$ | $N_r^{*,A}$ | $N_r^{*,B}$ |

Suppose 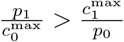. Table S2 shows an overview of the equilibrium ratios and their transitions for this case. Now, 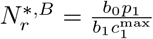 is without contradiction for 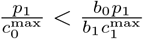. Violating this condition removes the contradiction for 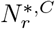

For 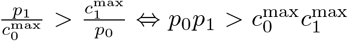, exactly one resource equilibrium will not exist regardless of the biomass conversion factors *b*_*i*_, whereas the other cases allow for existence of both equilibria for some paramter combinations. This phenomenon occurs because here, the overall resource production capabilities *p*_0_*p*_1_ overwhelm the maximum consumption capabilities 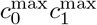 of both populations.

**Table S2.** Viability of the resource equilibria and contradictionless equilibrium ratios for 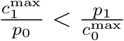.

| | $N_r^* < \frac{c_1^{\max}}{p_0}$ | $\frac{c_1^{\max}}{p_0} < N_r^* < \frac{p_1}{c_0^{\max}}$ | $\frac{p_1}{c_0^{\max}} < N_r^*$ |
| --- | --- | --- | --- |
| $R_0^*$ | does not exist | does not exist | exists |
| $R_1^*$ | exists | does not exist | does not exist |
| equilibrium ratio $N_r^*$ | $N_r^{*,C}$ | None | $N_r^{*,B}$ |

## SI 3 Cost-benefit model

As seen in Figure 3, the cost-benefit model (Equation 7) displays bistability between the trivial zero-equilibirum and an equilibrium ratio. Similar to the explicit resource exchange model (Equation 4), using resource consumption functions *c*_*i*_(*R*_*i*_) with upper limits 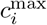 leads to three possible equilibrium ratio formulas. Which one will be assumed depends on the parameters. The proof for this is analogous to the one presented in Section SI 2.

Consider the system (*N*_*r*_, *R*_0_, *R*_1_) for *N*_0_ *>* 0, *N*_1_ *>* 0, *R*_0_ ≥ 0, *R*_1_ ≥ 0:

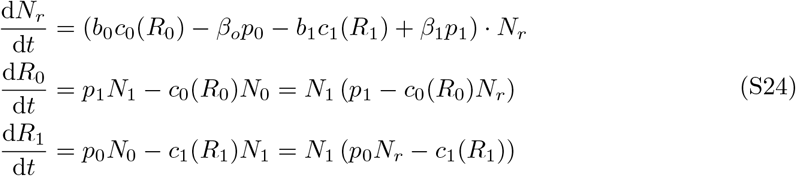

There is only one possible equilibrium under the assumption that the resource equilibria exist:

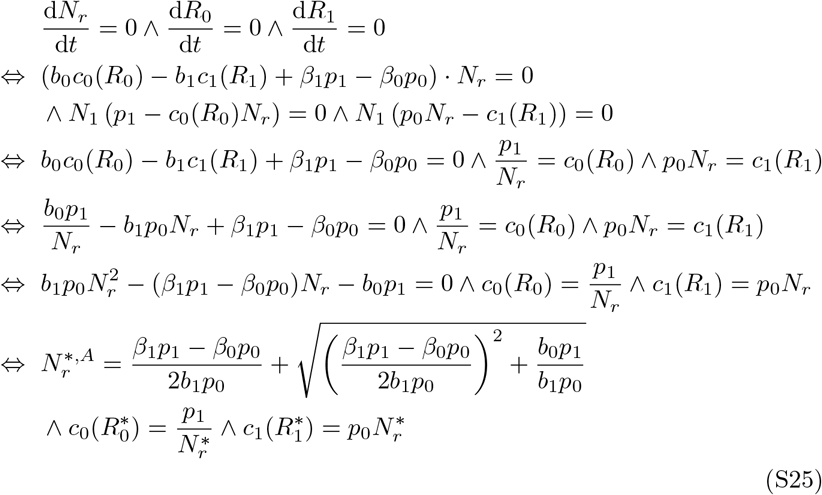

Note that the last step of this proof involves solving a quadratic equation which yields two possible signs before the square root in the formula for 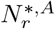. Since both populations and therefore the ratio are assumed to be strictly positive, only a positive sign is biologically relevant.

The growth rate at this ratio equilibrium is

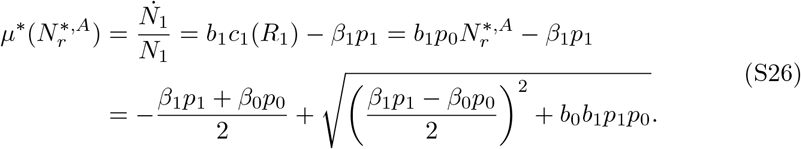

The Jacobian of the system is

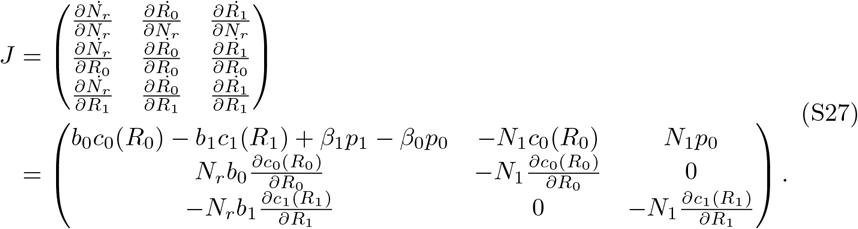

Evaluating *J* at the equilibrium point yields

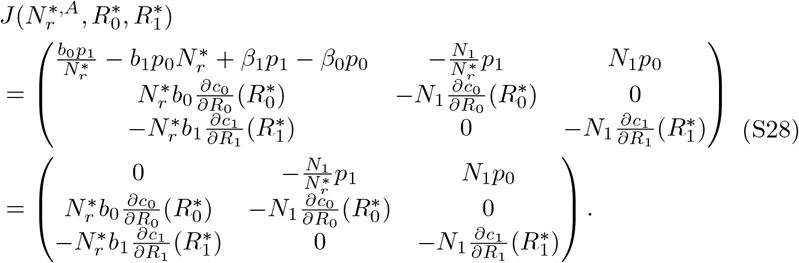

The remainder of this proof is exactly as in section SI 2, therefore 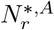 is asymptotically stable for the cost-benefit model as well.

The other two possible equilibrium ratios can be found by considering the cases where the resource uptake capabilities are saturated. If both resources are highly abundant, the system will always reach a population ratio which takes one of the resources below its maximum saturation level. We consider the case where the partial equilibrium for *R*_0_ exists and *R*_1_ grows exponentially alongside both populations (the opposite case is analogous). Then, *R*_0_ will eventually reach its equilibrium where *p*_1_*N*_1_ = *c*_0_(*R*_0_)*N*_0_ holds. *R*_1_ will increase, which eventually leads to 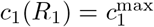. Applying this to the population ODEs yields:

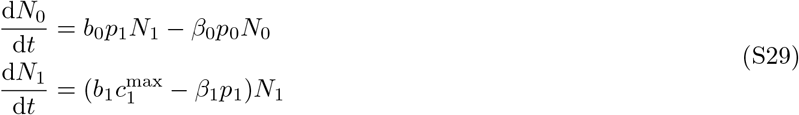

As discussed for Equation (S16), the steady states for all three modes of system behaviour exist for parameter values below some critical thresholds, in particular 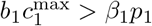. Therefore, *N*_1_ grows exponentially with rate

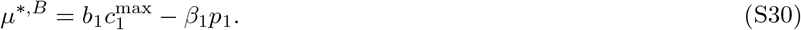

Evaluating the population ratio ODE for its equilibrium yields

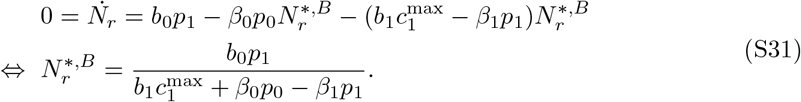

Analogously, we find

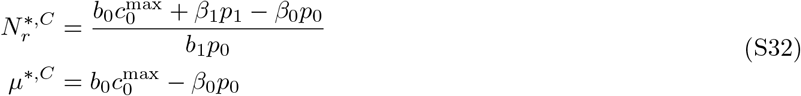

for the case where *R*_1_ stabilizes and *R*_0_ → ∞.

The three cases can again be differentiated analytically by calculating all three values for 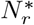, inserting them alongside 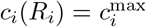 into the differential equations of the resources and checking which of the three formulas results in the signs of the resource differential equations matching the respective assumptions regarding the stability of the partial resource equilibria (compare section SI 2.2).

## SI 4 Dilution and external resource limitations

### SI 4.1 Chemostat without external resource

We modify the cost-benefit model (see Eq. 7) by including a constant dilution rate *γ*, resulting the the equations

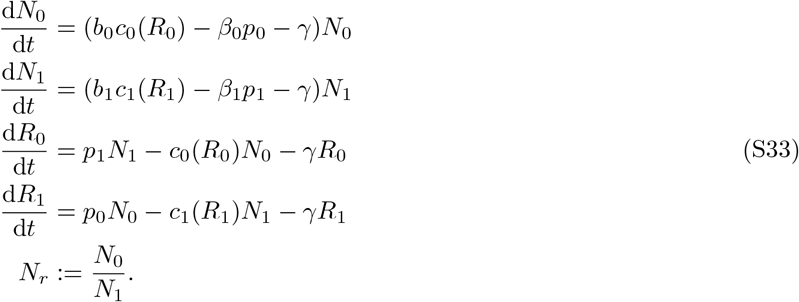

This simplified chemostat model has a trivial equilibrium (*N*_*i*_ = *R*_*i*_ = 0). The Jacobian at this equilibrium is

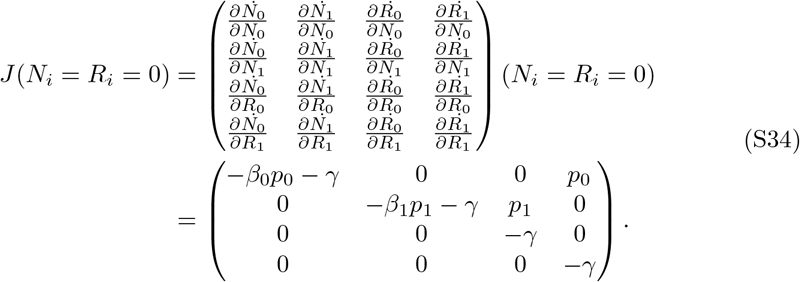

Since *J* is an upper triangular matrix, the eigenvalues are its diagonal elements, which are all strictly negative. Thus, the trivial equilibrium is always stable for *γ >* 0.

Figure S1 shows steady-state population ratios and growth rates for different dilution rates and initial conditions. The populations go extinct if the initial resource levels are too low. This minimum requirement rises with increasing *γ*. There exists a threshold for *γ* beyond which no initial conditions can lead to growth. The long-term population ratios are the same regardless of *γ* and the choice of initial conditions, as long as the populations do not go extinct. The sum of the long-term population growth rate and *γ* always reaches the same value.

### SI 4.2 External resource without dilution – the batch model

We simulate a typical batch experiment by employing Equation (11) and setting *γ* = 0. The dynamics of this system is exemplified in Figure S2

## SI 5 Forced crossfeeding in *E. coli* knockout mutant cocultures using dynamic FBA

Fig S3 shows simulations of an obligatory ΔI-ΔK cross-feeding system of two amino acid auxotroph *E. coli* populations. Exactly as for Fig 6, the qualitative and quantitative behaviour matches predictions from the explicit resource exchange model (Equation 4). After a transitional phase, both populations grow exponentially with the same rate and a constant ratio *N*_0_*/N*_1_ whereas the amino acid amounts stabilize.

Figure S4 shows the simulation for the ΔI-ΔT cross-feeding pair of *E. coli* auxotrophs. Here, the qualitative behaviour matches the explicit resource exchange model (Equation 4) and the simulations presented in Figures 6 and S3. However, the quantitative behaviour differs because the core assumption that the production pathways of both resources are independent, is violated. Threonine is a precursor to isoleusine. Therefore, the amount of isoleucine produced by the *Delta*T knockout strain is dependent on its threonine supply. For this, the custom formula predicting the equilibrium ratio

**Figure S2.**
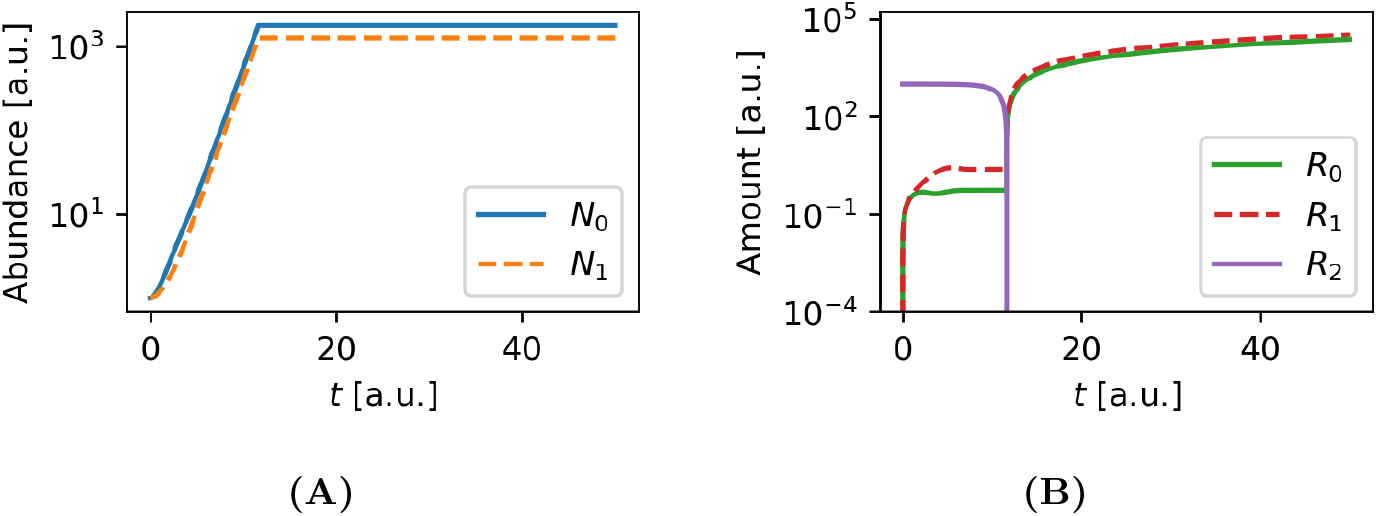
Approximation of exponential growth dynamics in batch cultures. The external resource model (Equation 11) without dilution (*γ* = 0) was used to simulate batch culture conditions. The Monod consumption functions with upper limits were used (Equation 10). The parameters were set to 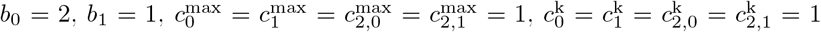 *β*_0_ = *β*_1_ = *γ* = *e* = 0, *p*_0_ = *p*_1_ = 0.5, *s*_0,2_ = 1 and *s*_1,2_ = 10. **(A)** Population abundances over time. First, there is an exponential growth phase where both populations converge towards a stable ratio and grow with the same relative rate. Then, both populations stop growing. **(B)** Resource amounts over time from the simulation used for (A). The exchanged resources *R*_0_ and *R*_1_ stabilize during the exponential growth phase of the populations. This phase ends when the external resource *R*_2_ becomes growth limiting. It is depleted to near zero quickly. After that, the other two resources accumulate during the time frame where the populations cease to grow.

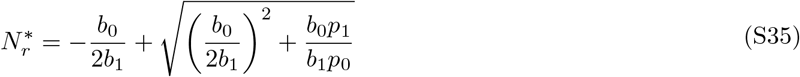

can be derived. Table S3 shows descriptions of the state variables and parameters of the ΔI-ΔI coculture used for deriving the formula.

At the equilibrium ratio, the per-capita growth rates of both populations have to be equal. Additionally, the resource production and consumption terms need to be equal for each resource.

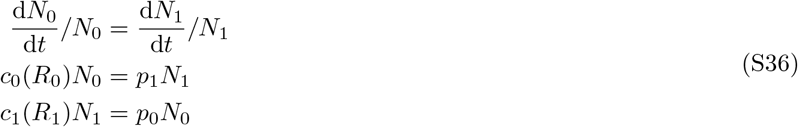

**Figure S3.**
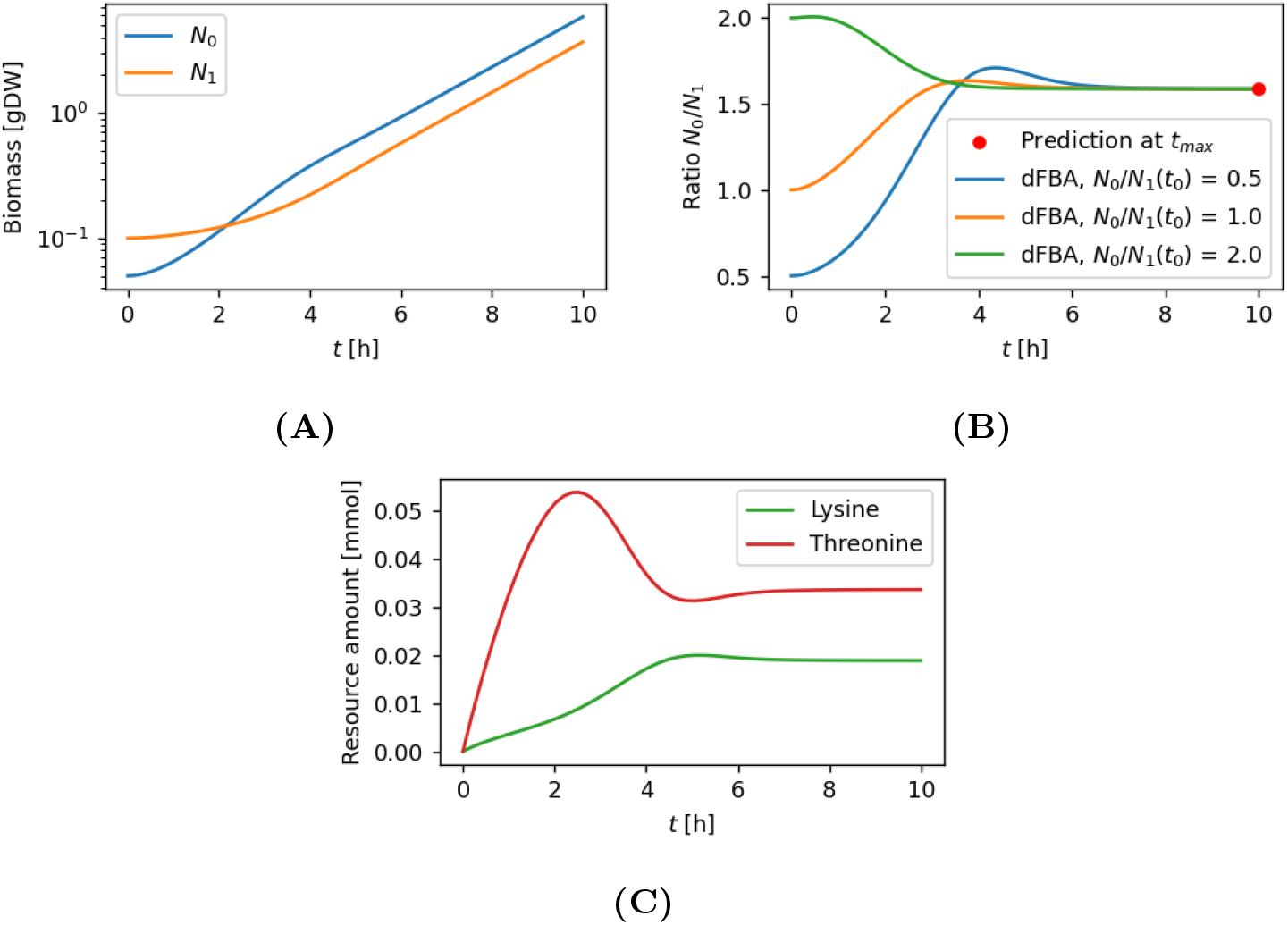
dFBA simulation of two *E. coli* populations lacking either lysine or threonine production capabilities. *N*_0_ is the lysine producer, *N*_1_ the threonine producer. The amino acid excretion rates were both set to 0.1 mmol/[gDW*h]. The uptake functions are given in Equation (10). The parameters are set to 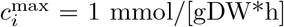 and 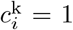 mmol. Both populations can grow due to exchange of the amino acids. **(A)** Population abundances over time. After a transitional phase, both populations grow exponentially with the same rate and a constant ratio *N*_0_*/N*_1_. **(B)** Predicted final population ratio and simulated ratios over time for different starting ratios. After a lag phase, the simulated ratios all stabilize at the same level. The prediction was calculated via the equilibrium ratio Equation (5) for the explicit resource exchange model (4). Simulation and prediction are in agreement. **(C)** Amino acid amounts over time for the simulation shown in (A). After the transitional phase, both resource levels stabilize.

**Figure S4.**
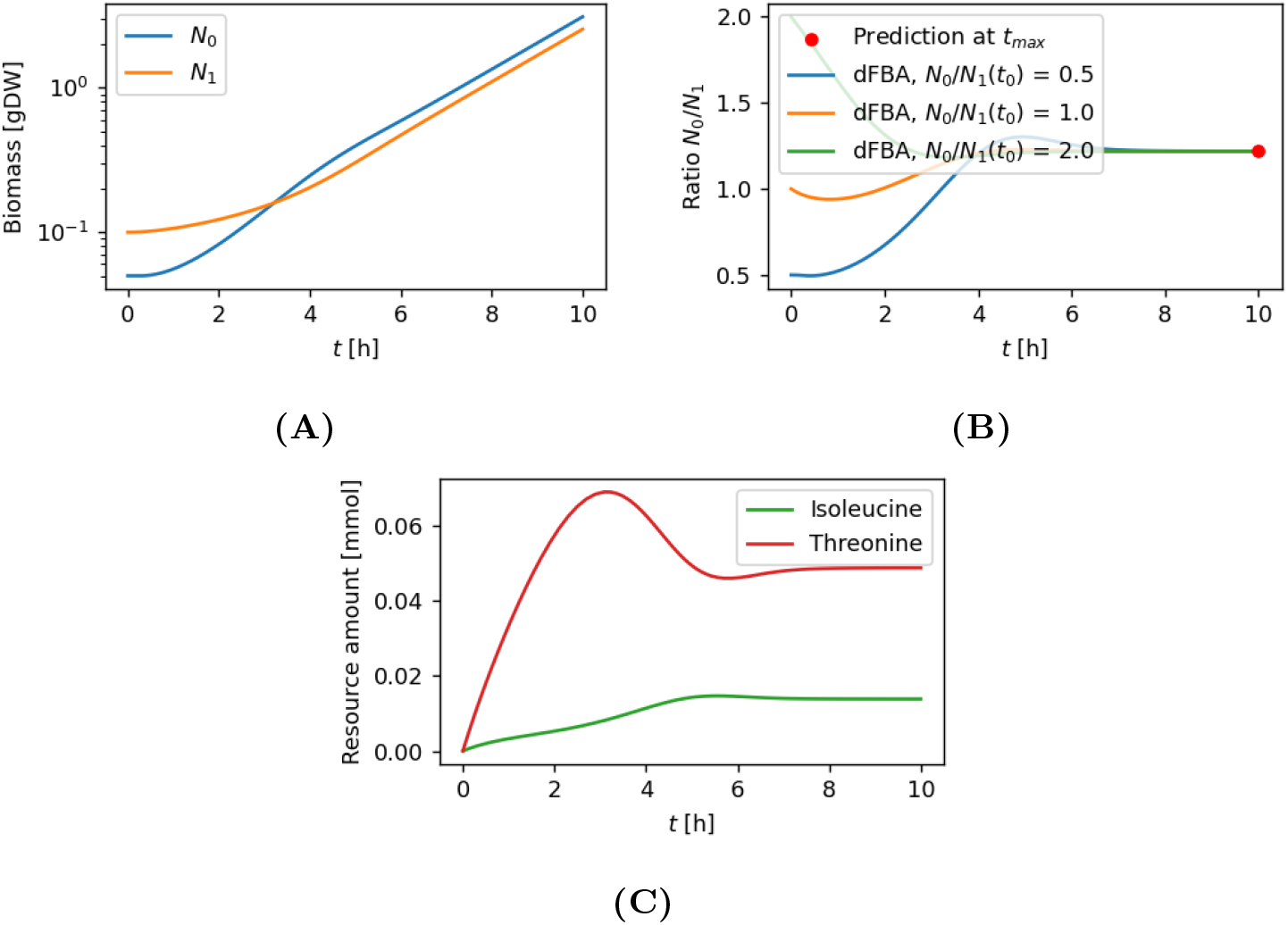
dFBA simulation of two *E. coli* populations lacking either isoleucine or threonine production capabilities. *N*_0_ is the isoleucine producer, *N*_1_ the threonine producer. The amino acid excretion rates were both set to 0.1 mmol/[gDW*h]. The uptake functions are given in Equation (10). The parameters are set to 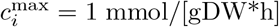 and 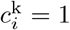 mmol. Both populations can grow due to exchange of the amino acids. **(A)** Population abundances over time. After a transitional phase, both populations grow exponentially with the same rate and a constant ratio *N*_0_*/N*_1_. **(B)** Predicted final population ratio and simulated ratios over time for different starting ratios. After a lag phase, the simulated ratios all stabilize at the same level. The prediction was calculated via the equilibrium ratio equation (S35) which was derived specifically for this scenario. Simulation and prediction are in agreement. **(C)** Amino acid amounts over time for the simulation shown in (A). After the transitional phase, both resource levels stabilize.

**Table S3.** State variables and parameters of the isoleucine-threonine knockout coculture.

| Variable or parameter | description |
| --- | --- |
| $N_1$ | THR consumer |
| $N_1$ | ILE consumer |
| $R_0$ | threonine (THR) |
| $R_1$ | isoleucine (ILE) |
| $b_0$ | effective THR into biomass conversion factor |
| $b_1$ | ILE into biomass conversion factor |
| $b_{\text{THR}}$ | THR into biomass conversion factor |
| $p_0$ | ILE production rate |
| $p_1$ | THR production rate |

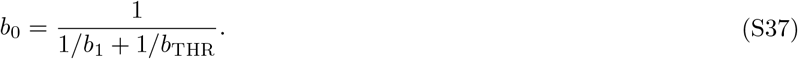

The effective biomass conversion factor for the threonine consumer is 1

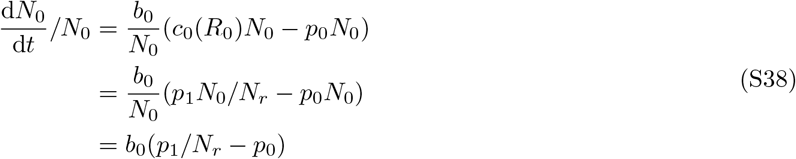

This yields a per-capita growth rate of the threonine consumer that is the biomass conversion factor multiplied with the difference of the threonine taken up and the threonine that is converted into isoleucine for excretion (the conversion rate is 1-to-1):

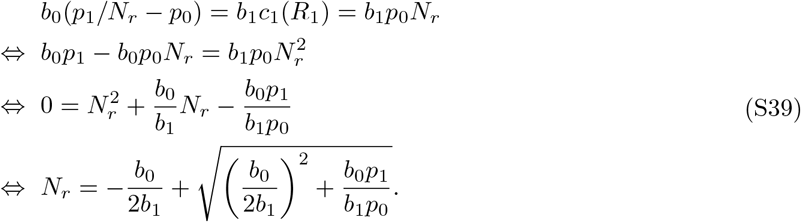

Setting both growth rates equal yields the formula for 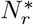 (note that the sign in front of the square root must be positive since the ratio has to be positive):

## References

[1] Erica C. Seth and Michiko E. Taga. “Nutrient cross-feeding in the microbial world”. In: Frontiers in Microbiology 5 (2014). ISSN: 1664-302X. DOI: 10.3389/fmicb.2014.00350.

[2] Jake N Barber, Aysha L Sezmis, Laura C Woods, Trenton D Anderson, Jasmyn M Voss, and Michael J McDonald. “The evolution of coexistence from competition in experimental co-cultures of Escherichia coli and Saccharomyces cerevisiae”. In: The ISME Journal 15.3 (Oct. 2020), pp. 746–761. ISSN: 1751-7370. DOI: 10.1038/s41396-020-00810-z.

[3] Elena Kazamia, Hjördis Czesnick, Thi Thanh Van Nguyen, Martin Tom Croft, Emma Sherwood, Severin Sasso, Sarah James Hodson, Martin James Warren, and Alison Gail Smith. “Mutualistic interactions between vitamin B12-dependent algae and heterotrophic bacteria exhibit regulation”. In: Environmental Microbiology 14.6 (Mar. 2012), pp. 1466–1476. ISSN: 1462-2920. DOI: 10.1111/j.1462-2920.2012.02733.x.

[4] Matthew AA Grant, Elena Kazamia, Pietro Cicuta, and Alison G Smith. “Direct exchange of vitamin B12 is demonstrated by modelling the growth dynamics of algal–bacterial cocultures”. In: The ISME Journal 8.7 (Feb. 2014), pp. 1418–1427. DOI: 10.1038/ismej.2014.9.

[5] Sander Sieuwerts. “Microbial Interactions in the Yoghurt Consortium: Current Status and Product Implications”. In: SOJ Microbiology & Infectious Diseases 4.2 (2016), pp. 01–05. ISSN: 2372-0956. DOI: 10.15226/sojmid/4/2/00150.

[6] Freddy Bunbury, Evelyne Deery, Andrew P. Sayer, Vaibhav Bhardwaj, Ellen L. Harrison, Martin J. Warren, and Alison G. Smith. “Exploring the onset of B12-based mutualisms using a recently evolved Chlamydomonas auxotroph and B12-producing bacteria”. In: Environmental Microbiology 24.7 (May 2022), pp. 3134–3147. ISSN: 1462-2920. DOI: 10.1111/1462-2920.16035.

[7] Jan Dolinšek, Josep Ramoneda, and David R Johnson. “Initial community composition determines the long-term dynamics of a microbial cross-feeding interaction by modulating niche availability”. In: ISME Communications 2.1 (Aug. 2022). ISSN: 2730-6151. DOI: 10.1038/s43705-022-00160-1.

[8] Martina Du, Jeremy M. Chácon, Heejoon Park, Campbell Putnam, Tomáš Gedeon, William R. Harcombe, and Ross P. Carlson. “Cellular Economics of Exchanged Metabolites Alter Ratios of Microbial Trading Partners in a Predictable Manner”. In: ACS Synthetic Biology 14.11 (2025). PMID: 41071854, pp. 4373–4387. DOI: 10.1021/acssynbio.5c00265. eprint: https://doi.org/10.1021/acssynbio.5c00265.

[9] Andreu Mas-Colell, Michael Dennis Whinston, Jerry R Green, et al. Microeconomic theory. Vol. 1. Oxford university press New York, 1995.

[10] Joshua Tasoff, Michael T. Mee, and Harris H. Wang. “An Economic Framework of Microbial Trade”. In: PLoS ONE (2015). DOI: 10.1038/msb.2011.65.

[11] J. J. Heijnen, M. C. M. van Loosdrecht, and L. Tijhuis. “A black box mathematical model to calculate auto- and heterotrophic biomass yields based on Gibbs energy dissipation”. In: Biotechnology and Bioengineering 40.10 (Dec. 1992), pp. 1139–1154. DOI: 10.1002/bit.260401003.

[12] Anne Goelzer and Vincent Fromion. “Resource allocation in living organisms”. In: Biochemical Society Transactions 45.4 (July 2017), pp. 945–952. DOI: 10.1042/bst20160436.

[13] Hugo Dourado, Wolfram Liebermeister, Oliver Ebenhöh, and Martin J. Lercher. “Mathematical properties of optimal fluxes in cellular reaction networks at balanced growth”. In: PLOS Computational Biology 19.6 (June 2023). Ed. by Pedro Mendes, e1011156. DOI: 10.1371/journal.pcbi.1011156.

[14] Chen Liao, Tong Wang, Sergei Maslov, and Joao B. Xavier. “Modeling microbial cross-feeding at intermediate scale portrays community dynamics and species coexistence”. In: PLOS Computational Biology 16.8 (Aug. 2020). Ed. by Jacopo Grilli, e1008135. ISSN: 1553-7358. DOI: 10.1371/journal.pcbi.1008135.

[15] Jeffrey D. Orth, Tom M. Conrad, Jessica Na, Joshua A. Lerman, Hojung Nam, Adam M. Feist, and Bernhard Ø Palsson. “A comprehensive genome-scale reconstruction of Escherichia coli metabolism—2011”. In: Molecular Systems Biology 7.1 (2011).

[16] Radhakrishnan Mahadevan, Jeremy S Edwards, and Francis J Doyle. “Dynamic flux balance analysis of diauxic growth in Escherichia coli.” eng. In: Biophys J 83.3 (Sept. 2002), pp. 1331–1340. DOI: 10.1016/S0006-3495(02)73903-9. URL: http://dx.doi.org/10.1016/S0006-3495(02)73903-9.

[17] Ali Ebrahim, Joshua A. Lerman, Bernhard Ø. Palsson, and Daniel R. Hyduke. “COBRApy: COnstraints-Based Reconstruction and Analysis for Python”. In: BMC Systems Biology (2013). DOI: 10.1186/1752-0509-7-74.

[18] Tyler D. Ross, Hanhyeok Im, Brennan G. Keogh, Christopher A. Klausmeier, and Ophelia S. Venturelli. “Metabolic interplay drives population cycles in a cross-feeding microbial community”. In: Nature Communications 16.1 (Oct. 2025). ISSN: 2041-1723. DOI: 10.1038/s41467-025-63986-y.

[19] Zarath M Summers, Heather E Fogarty, Ching Leang, Ashley E Franks, Nikhil S Malvankar, and Derek R Lovley. “Direct exchange of electrons within aggregates of an evolved syntrophic coculture of anaerobic bacteria”. In: Science 330.6009 (2010), pp. 1413–1415.

